# Cell-Type-Specific Epigenetic Priming of Gene Expression in Nucleus Accumbens by Cocaine

**DOI:** 10.1101/2022.06.24.497533

**Authors:** Philipp Mews, Yentl Van der Zee, Hope Kronman, Ashik Gurung, Aarthi Ramakrishnan, Caleb Browne, Rita Futamura, Molly Estill, Meagan Ryan, Abner A Reyes, Benjamin A Garcia, Simone Sidoli, Li Shen, Eric J Nestler

**Affiliations:** Nash Family Department of Neuroscience and Friedman Brain Institute, Icahn School of Medicine at Mount Sinai, New York, NY; Department of Biochemistry and Molecular Biophysics, Washington University School of Medicine, St. Louis, MO; Department of Biochemistry, Albert Einstein College of Medicine, New York, NY

**Keywords:** withdrawal, relapse, epigenetics, gene priming, chromatin remodeling

## Abstract

A hallmark of addiction is the ability of drugs of abuse to trigger relapse after periods of prolonged abstinence. Here, we describe a novel epigenetic mechanism whereby chronic cocaine exposure causes lasting chromatin and downstream transcriptional modifications in the nucleus accumbens (NAc), a critical brain region controlling motivation. We link prolonged withdrawal from cocaine to the depletion of the histone variant H2A.Z, coupled to increased genome accessibility and latent priming of gene transcription, in D1 dopamine receptor-expressing medium spiny neurons (D1 MSNs) that relates to aberrant gene expression upon drug relapse. The histone chaperone ANP32E removes H2A.Z from chromatin, and we demonstrate that D1 MSN-selective *Anp32e* knockdown prevents cocaine-induced H2A.Z depletion and blocks cocaine’s rewarding actions. By contrast, very different effects of cocaine exposure, withdrawal, and relapse were found for D2-MSNs. These findings establish histone variant exchange as an important mechanism and clinical target engaged by drugs of abuse to corrupt brain function and behavior.

## INTRODUCTION

Chronic cocaine exposure induces persistent changes in gene regulation in the brain’s motivation and reward circuitry, coupled to neuroplasticity within the brain and vulnerability to relapse. Susceptibility to relapse is believed to involve in part stable changes in chromatin in the nucleus accumbens (NAc), a brain region that controls motivated behaviors, that alter transcription during long-term drug withdrawal (*1*). However, the molecular events that underlie maladaptive gene activity in NAc in cocaine addiction and other substance use disorders remain incompletely understood. Here, we investigated how withdrawal from chronic exposure to cocaine changes gene regulation and the underlying chromatin landscape in defined cell populations within the NAc and causally linked these changes to cocaine-associated behavioral phenotypes.

The NAc is primarily composed of two opposing types of GABAergic projection neurons called medium spiny neurons (MSNs), expressing either D1 dopamine receptors (DRD1) or D2 dopamine receptors (DRD2). These distinct subtypes exhibit differences in cocaine-related neural activity and effects on behaviors associated with drug exposure: activation of D1 MSNs promotes reward-associated behaviors, whereas activation of D2 MSNs suppresses them (*2–10*). Recent studied showed that acute cocaine experience induces immediate-early gene expression in D1 MSNs, and that the transcription factors CREB and AP-1 drive dopamine-related gene expression programs in the NAc (*11–14*).

While these data are interesting, they do not explain the long-lasting nature of cocaine-related changes – which have been shown to be persistent months following the last cocaine exposure. Here, we show that prolonged drug withdrawal is characterized by latent changes in gene regulation in D1 MSNs that become apparent upon cocaine relapse, and that this transcriptional priming is associated with persistent remodeling of chromatin structure in this cell type. Recent work by our and other laboratories has characterized cocaine-induced changes in histone modifications (*15–22*), which play critical roles in orchestrating stimulus-related gene activity (*16, 23–25*). However, little is known about how epigenetic changes relate to lasting changes in gene regulation in withdrawal and relapse: prior studies have failed to demonstrate appreciable overlap of any given histone modification to changes in gene expression.

One explanation for such relatively limited overlap is the possibility that the field has been focusing on the wrong histone modifications: studying those histone modifications that are best characterized in other systems (mostly other tissues) as opposed to using an open-ended approach to first identify the histone modifications that are most robustly affected in NAc MSNs in response to cocaine exposure. To overcome this gap in knowledge, we took advantage of unbiased mass spectrometry to characterize in a comprehensive manner withdrawal-associated changes in both histone modifications and the abundance of histone variants, which are emerging as important regulators of gene expression in the adult nervous system (*26, 27*). Histone variants are non-allelic counterparts of canonical histones that are structurally and functionally distinct, with replication-independent dynamics that are controlled by a network of histone chaperones in post-mitotic cells (*28*). We found that prolonged withdrawal from cocaine causes marked depletion of the histone variant, H2A.Z, especially at putative enhancer regions and key neuronal genes that control synaptic plasticity. We show further that genome accessibility is increased prominently at these genes after prolonged withdrawal, and linked to aberrant gene expression upon drug relapse, specifically in D1 MSNs. Removal of H2A.Z is dependent upon the histone chaperone ANP32E (*29*), and we demonstrate that D1 MSN-selective *Anp32e* knockdown prevents cocaine-induced H2A.Z depletion and effectively blocks cocaine conditioned place preference (CPP), an indirect measure of cocaine reward. By contrast, chronic cocaine exposure, prolonged withdrawal, and relapse produce very different patterns of transcriptional regulation in D2 MSNs. These findings thus identify highly novel circuit-specific epigenetic priming of gene expression in NAc as a critical mechanism of the lasting effects of cocaine on the brain.

### Cocaine challenge following drug withdrawal reveals latent dysregulation of D1 MSN transcription

A principal hallmark of addiction is the ability of drugs of abuse like cocaine to trigger relapse after periods of prolonged abstinence. An emerging research focus therefore are mechanisms by which neuronal gene regulation is primed (or desensitized) in the NAc through a course of prolonged withdrawal for aberrant expression upon drug relapse. Yet, little is known about how cocaine lastingly impacts circuit-specific gene programs in the divergent D1 and D2 MSNs of this brain region. Here, we first compared cell-type-specific transcriptional responses to the acute experience of cocaine in drug-naïve mice with the responses elicited by the equivalent dose of cocaine but after 30 d of withdrawal from chronic cocaine exposure (Fig. 1A). To map transcriptional activities in D1 vs. D2 MSNs, we employed two transgenic mouse models (*Drd1a* / *Drd2a*::EGFP-L10a) that enabled fluorescent-activated nuclei sorting (FANS) of these functionally-distinct cell types, combined with RNA-sequencing (RNA-seq) (Fig. 1B; fig. S1). Signatures of subtype-defining gene transcripts confirmed the selectivity of this technical approach across study groups (fig. S2A).

**Fig. 1.**
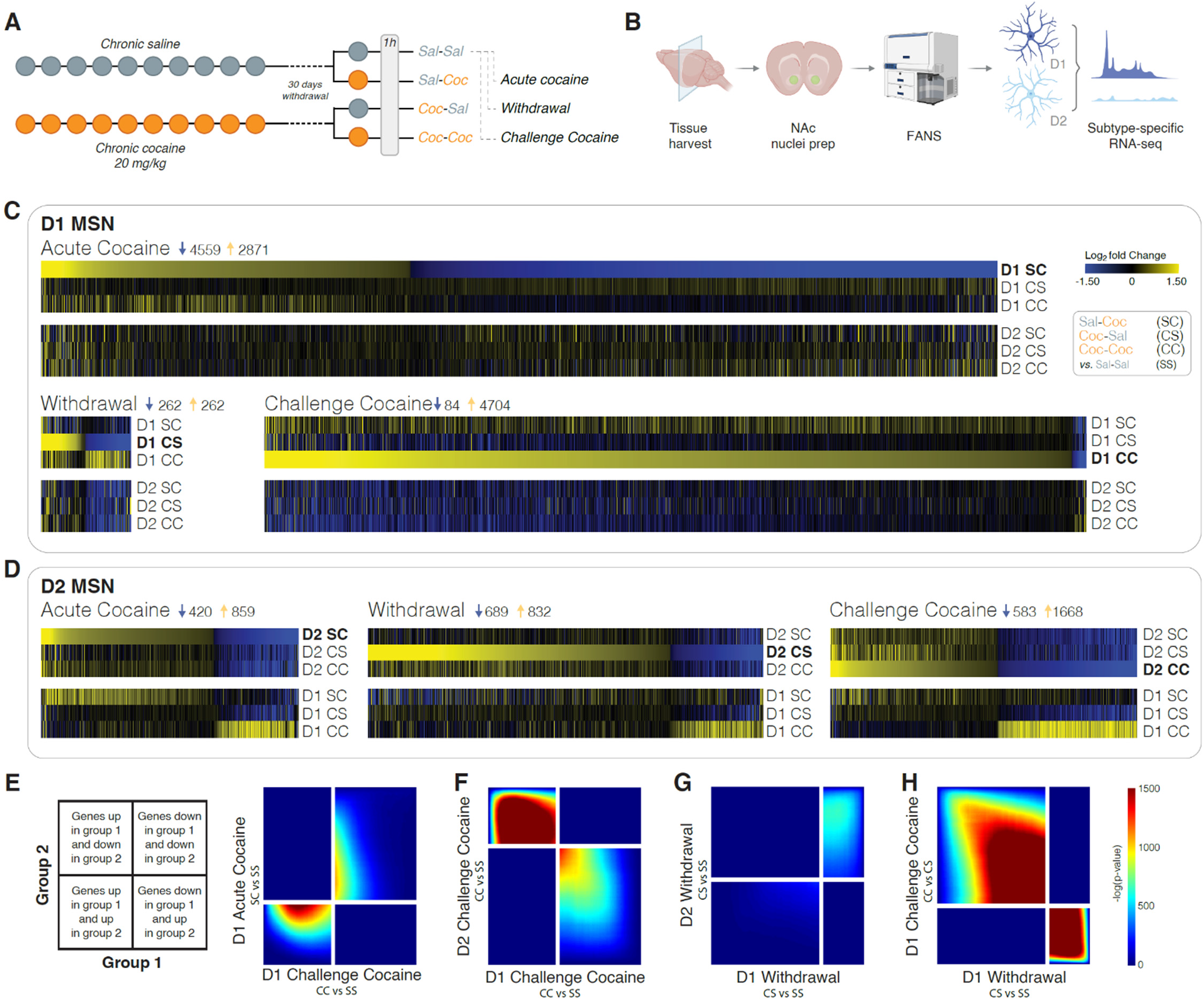
Latent dysregulation of D1 MSN gene transcription in NAc after prolonged withdrawal from chronic cocaine. (**A**) Experimental outline. To characterize cell-type-specific changes in gene expression following prolonged cocaine withdrawal, we employed two transgenic mice (*Drd1a* / *Drd2a*::EGFP-L10a) for purification of D1 and D2 MSN nuclei. Following withdrawal (30 d) from chronic saline or cocaine (10 d), animals received either a saline or cocaine (20 mg/kg challenge injection *i.p.*) in order to profile and compare the transcriptional response in D1 vs. D2 MSNs upon acute cocaine in drug-naïve mice (SC, Saline-Cocaine) with that to a cocaine challenge in withdrawal animals (CC, Cocaine-Cocaine), and to a saline challenge in withdrawal animals (CS, Cocaine-Saline), when compared to control saline animals (SS, Saline-Saline). (**B**) Nuclei were prepared from NAc of individual mice analyzed 1 h after the last challenge injection, followed by FANS for MSN subtype-specific RNA-seq (6-8 animals per group). (**C, D**) Heatmaps show genes differentially expressed in **(C)** D1 MSNs and **(D)** D2 MSNs following acute cocaine (SC vs. SS), withdrawal (CS vs. SS), and cocaine challenge (CC vs. SS), across all treatment groups, revealing latent changes in gene regulation during withdrawal primarily in D1 MSNs. (**E-H**) Comparison of transcriptome-wide expression profiles in a threshold-free manner using rank-rank hypergeometric overlap (RRHO, key to the left for interpretation), with correction for multiple comparisons using the Benjamini–Hochberg procedure. **(E)** RRHO comparing transcriptional changes in D1 MSNs after acute cocaine (SC) and challenge cocaine (CC) confirmed partial directional overlap. In contrast, **(F)** opposite regulation of gene expression was evident in D2 MSNs compared to D1 MSNs upon a cocaine challenge after withdrawal (CC), modelling cocaine relapse. **(G)** RRHO comparing transcriptional changes in D1 and D2 MSNs after cocaine withdrawal (CS) showed no overlap. **(H)** Threshold-free comparison of transcriptomes in D1 MSNs indicates that a cocaine challenge (CC) largely reverses changes in gene expression linked to withdrawal (CS).

We initially evaluated how an acute dose of cocaine in drug-naïve mice (SC, Sal-Coc) changes the D1 and D2 transcriptomes of the NAc. Compared to D2 MSNs, D1 MSNs showed a pronounced transcriptional response to cocaine involving gene programs related to neural plasticity, such as dendritic spine and post-synaptic density (fig. S3). In animals with a distant history of chronic cocaine, drug relapse after prolonged withdrawal (CC, Coc-Coc) stimulated gene expression to a substantially greater extent than any other condition, prompting an amplified transcriptional response in D1 MSNs (Fig. 1C). In contrast, cocaine-related gene expression upon relapse remained similar in D2 MSNs when compared to acute cocaine in drug-naïve mice (CC and SC, respectively; Fig. 1D). Notably, withdrawal-related dysregulation of D1 MSN-specific gene expression is latent: the D1 transcriptome remained largely unchanged in withdrawal animals with a priming dose of saline (CS, Coc-Sal) compared to baseline controls (SS, Sal-Sal; Fig. 1C). By contrast, upon a relapse dose of cocaine, many immediate-early genes were primed for rapid induction, and numerous other genes, previously not affected by cocaine in drug-naïve mice, were “primed” during withdrawal and induced by a cocaine challenge (CC, fig. S2B-C). To evaluate the transcriptome-wide degree of overlap between the differential gene expression profiles upon acute cocaine (SC) vs. drug challenge (CC), we performed threshold-free rank-rank hypergeometric overlap (RRHO) analysis. This test showed that the impact of cocaine relapse on the D1 transcriptome, albeit dramatically widened and strengthened, corresponded directionally to the effects of its initial acute experience (Fig. 1E). Compared to D1 MSNs, very different patterns were evident for D2 MSNs, including sets of genes that changed in the opposite direction compared to D1 MSNs (Fig. 1F, G). Withdrawal-related changes in D1 MSN gene expression were largely reversed by cocaine challenge (Fig. 1H).

To explore what biological processes and cellular signaling pathways are affected by cocaine relapse (CC), we performed gene ontology (GO) analyses, which revealed that genes stimulated in D1 MSNs execute not only postsynaptic and somatodendritic functions but also have ribosomal and mitochondrial roles (fig. S4A). In contrast, D2 MSNs displayed downregulation of ribosomal subunit and mitochondrial proteins upon relapse (Fig. 1F, fig. S4B). We explored the upstream transcription factors that orchestrate relapse-induced gene programs to deduce regulatory players recruited in D1 MSNs by cocaine exposure, withdrawal, and relapse. Ingenuity Pathway Analysis (IPA) indicates a causal network of BDNF, EGF, and CAMK1 as upstream regulators, i.e., factors that have been previously implicated in dopamine- and cocaine-related gene transcription within the NAc (fig. S5A) (*30–32*). Moreover, levodopa and cocaine were predicted as the top upstream chemical activators, highlighting that our subtype-specific transcriptome analyses reflect well-documented regulation of D1 MSNs by cocaine (fig. S5B) (*6, 10, 33*). Given that these findings reveal latent changes in gene regulation in D1 MSNs after prolonged withdrawal prior to the relapse dose of cocaine, we set out to explore whether alterations in chromatin structure may underlie this phenomenon.

### Chronic cocaine causes lasting changes in D1 MSN chromatin structure

To investigate whether repeated cocaine experience lastingly alters subtype-specific chromatin landscapes, we examined ‘open’ chromatin regions in both D1 and D2 MSN nuclei genome-wide by use of assay for transposase-accessible chromatin with sequencing (ATAC-seq) (Fig. 2A). At baseline (SS), genic chromatin around transcriptional start sites (TSSs) was overall less accessible in D1 MSNs vs. D2 MSNs, which corroborates previous findings of higher D2 transcriptome diversity (*34*). In contrast, acute cocaine (SC) was linked to a dramatic genome-wide ‘opening’ of chromatin in D1 MSNs, with a far smaller effect apparent in D2 MSNs. Following chronic cocaine exposure and prolonged withdrawal (CS), D1-specific chromatin opening is sustained and, upon cocaine relapse (CC), linked to further globally increased accessibility, indicative of transcriptional priming within D1 MSNs (Fig. 2A). As before, changes in D2 MSNs were much smaller in magnitude at this genome-wide scale.

**Fig. 2.**
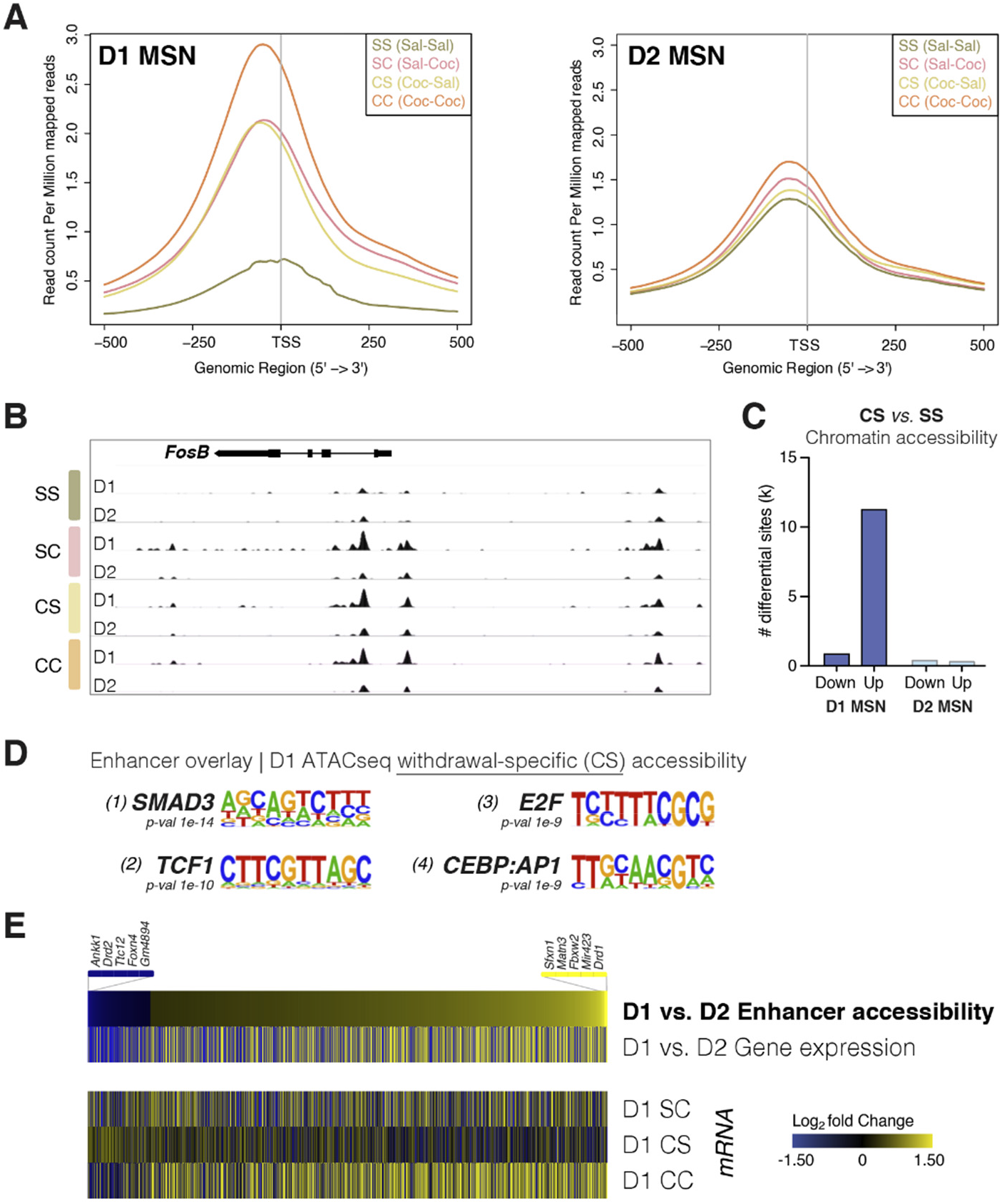
Withdrawal from chronic cocaine dramatically alters chromatin accessibility in D1 MSNs of NAc. (**A**) D1 and D2 MSN chromatin accessibility around TSSs was compared across treatment groups using ATAC-seq. SS (Saline-Saline, control saline), SC (Saline-Cocaine, acute cocaine in drug-naïve mice), CS (Cocaine-Saline, withdrawal from chronic cocaine), and CC (Cocaine-Cocaine, cocaine challenge following withdrawal from chronic cocaine) (see Fig. 1A). (**B**) Genome-browser view of the *Fosb* gene locus showing increased accessibility in D1 MSNs following cocaine exposure, both over its promoter and a putative enhancer region upstream. (**C**) Differentially accessible genomic loci following prolonged withdrawal from chronic cocaine (CS), showing predominant opening (Up) of chromatin in D1 MSNs. (**D**) Motif analysis on enhancer regions that showed increased accessibility specifically in D1 MSNs following withdrawal (Homer *de-novo* motif analysis). (**E**) Heatmap of differentially accessible enhancers between D1 and D2 MSNs, showing corresponding D1 vs. D2 MSN expression of the most proximal genes. The heatmap below shows corresponding gene expression in D1 MSNs with acute cocaine (SC), withdrawal (CS), and cocaine challenge (CC) compared to saline control (SS).

Early investigations of drug regulation of gene expression showed that cocaine upregulates ΔFOSB, encoded by *Fosb—*a member of the activator protein-1 (AP-1) family of transcription factors, specifically in D1 MSNs (*35, 36*). We found that, upon exposure to acute cocaine, both the *Fosb* core promoter and its upstream enhancer sites displayed increased accessibility in D1 but not D2 MSNs (Fig. 2B). This opening is sustained during prolonged withdrawal and relapse. In fact, *Fosb* is the third-most primed gene in D1 MSNs.

Genome-wide examination of differential gene accessibility following withdrawal in both neuronal subtypes confirmed widespread chromatin opening, and far fewer sites of decreased accessibility, in D1 MSNs, with a relatively small number of changes observed in D2 MSNs (Fig. 2C). Gene ontology analysis revealed that gene cohorts involved in regulating glutamatergic transmission, postsynaptic density assembly, and AMPA receptor activity are more accessible following withdrawal in D1 MSNs compared to drug-naïve mice (fig. S6). These D1-specific findings corroborate prior studies on whole NAc, which linked changes in AMPA and glutamatergic transmission to cocaine-induced structural plasticity during withdrawal (*37–39*), along with studies that demonstrate distinct effects of cocaine on glutamatergic function in D1 vs. D2 MSNs (*8*). Notably, we found that numerous ionotropic receptors and ion channels, important for neuronal excitability and synaptic plasticity, exhibit transcriptional priming or desensitization following prolonged withdrawal from chronic cocaine, including many glutamate receptors and potassium channels in D1 MSNs (fig. S7).

We next investigated whether the gene priming induced in D1 MSNs involves increased enhancer accessibility following cocaine withdrawal. We developed a machine learning model that was trained on histone mark ChIP-sequencing (ChIP-seq) data from the ENCODE project. Although enhancers are known to be cell-type-specific (*40*), their chromatin signatures are remarkably stable across different cell types, thus enabling their genome-wide discovery from our previously published NAc histone mark ChIP-seq dataset (*15*). We identified 7,796 putative enhancer sites using DeepRegFinder (*41*) (fig. S8A). In D1 MSNs, these newly identified enhancers are marked by increased accessibility in withdrawal, potentially priming neuronal gene programs for rapid induction upon cocaine relapse (fig. S8B-C). To gain insight into the cellular signaling pathways and transcription factors facilitating such latent changes in regulated gene expression, we performed *de novo* transcription factor motif analysis over enhancer sites that become accessible in withdrawal (Fig. 2D). This approach revealed that the withdrawal-related class of enhancers in D1 MSNs is marked by SMAD3 and E2F transcription factor binding elements, as well as by heterodimers of CCAAT enhancer-binding protein (C/EBP) and AP-1, which can recruit co-activators to open chromatin structure (Fig 2D). Notably, activin-receptor signaling via SMAD3 has been shown to contribute to cocaine-induced transcriptional and morphological plasticity in rat NAc, and to be upregulated following withdrawal from chronic cocaine (*22*). Further, our previous whole tissue transcriptome studies across the mouse brain reward circuitry predicted the E2F family of transcription factors as upstream regulators of NAc gene programs whose altered expression upon chronic cocaine or cocaine withdrawal is predictive of addiction-like behavior (*42*). Indeed, E2F3a in NAc was directly implicated in controlling transcriptional and behavioral responses to chronic cocaine (*43*).

To confirm that chromatin remodeling of cocaine-responsive enhancers is linked to gene expression in the NAc, we examined transcriptional activity of nearby genes in both D1 and D2 MSNs. We confirmed that differential enhancer accessibility between these neuronal subtypes is linked to cell-type-specific changes in gene expression, with D1-specific enhancer openness linked to increased mRNA expression of the most proximal genes in D1 neurons, and *vice versa* (395 preferentially accessible D2 enhancers, 2878 preferential D1 enhancers in CC, Fig. 2E). Nearby gene expression in D1 neurons was markedly increased following cocaine relapse, compared to respective gene expression induced by acute cocaine or in withdrawal from chronic cocaine (Fig. 2E). Together, these data indicate that enhancers targeted by SMAD, E2F, and AP-1 –all known to interact with the cocaine-responsive chromatin remodeler BRG1 that regulates cocaine-seeking behavior (*22*)– show increased accessibility in D1 MSNs after prolonged cocaine withdrawal, and are linked to priming of neuronal genes for rapid induction upon drug relapse.

### Withdrawal from chronic cocaine is associated with marked depletion of H2A.Z

Epigenetic remodeling has emerged as a potent switch for enhancer activity in plasticity-related transcription, with histone protein modifications believed to control enhancer and promoter function in withdrawal and transcription upon drug relapse (*1*). To systematically characterize global changes in histone modifications across cocaine exposure, withdrawal, and relapse, we took an unbiased approach by utilizing quantitative mass spectrometry on histones purified from NAc nuclei (Fig. 3A, fig. S9A). We calculated the relative abundance of single and combinatorial histone marks on each peptide—as just one example for acetylation and methylation of histone H3 lysine 9 and 14 (H3K9K14), which is reflective of their proportional genomic distribution (fig. S9B). Both acute cocaine and prolonged withdrawal triggered changes in key gene-regulatory histone marks, including trimethylation of H3K9, which our laboratory has previously shown to be induced transiently by acute cocaine (*21*). Beyond histone modifications per se, recent studies implicate dynamic turnover and exchange of histone variants as epigenomic regulators of neural plasticity (*44*). We observed dramatic depletion of the replication-independent histone variant, H2A.Z, following withdrawal from chronic cocaine by mass spectrometry (Fig. 3B), which was substantiated by Western blotting (fig. S10). H2A.Z, encoded by the *H2afz* gene, has multiple roles in transcription during cellular differentiation and development, and stimulus-induced removal of this variant from chromatin was shown to facilitate long-term hippocampal memory, indicating a memory suppressor function in the adult brain (*45*). Notably, our RNA-seq dataset also revealed significant downregulation of *H2afz* mRNA expression following withdrawal in both D1 and D2 MSNs (D1 CS*vs*SS log-FC = −1.3, p = 0.01; D2 CS*vs*SS log-FC = −1.5, p = 0.01, Fig. 3C).

**Fig. 3.**
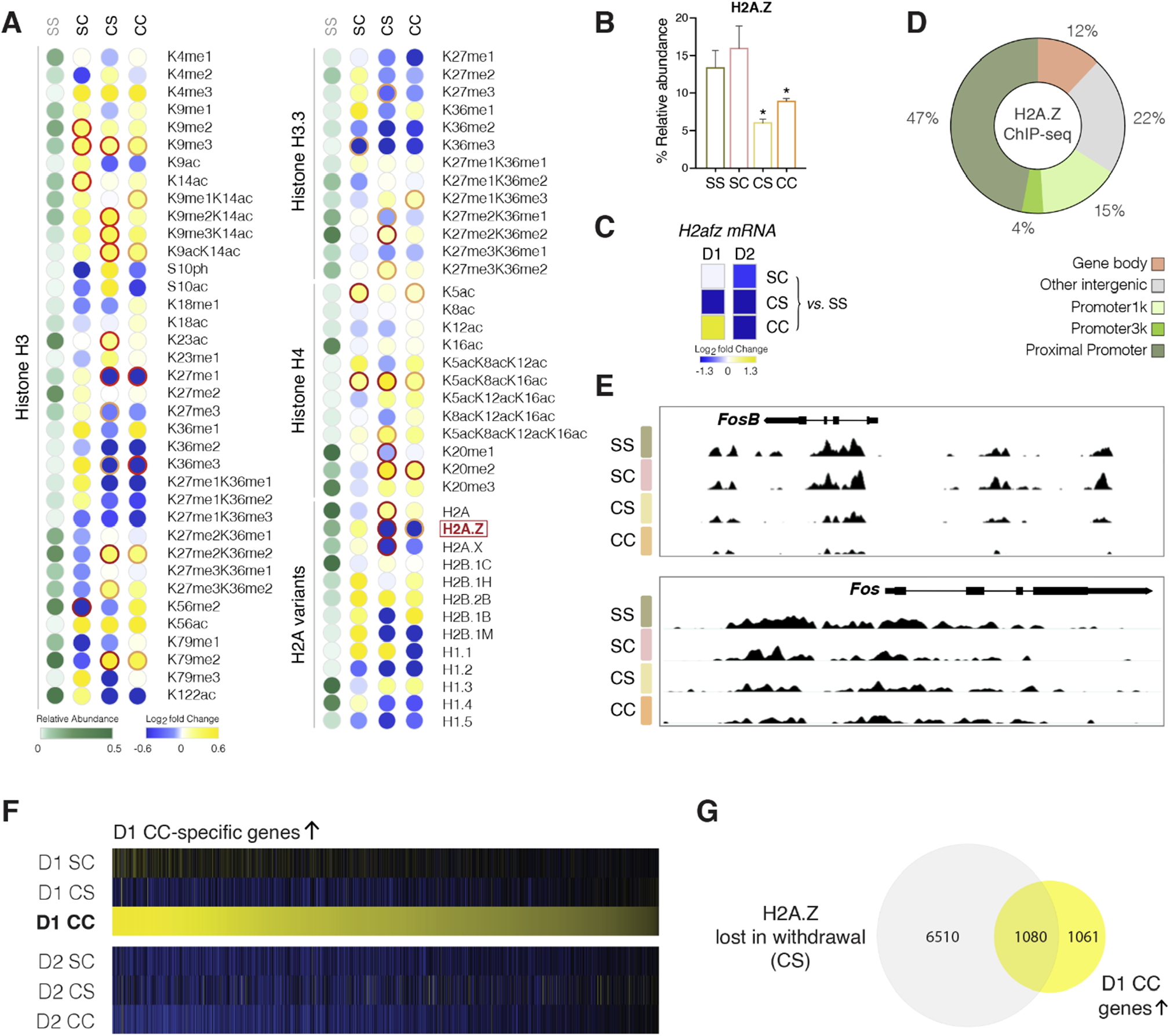
Mass spectrometry reveals cocaine-induced changes in post-translational histone modifications abundance of histone variants in NAc. (**A**) Relative abundance of each indicated modification on histone lysine (K) and serine (S) residues, or of each histone variant, is shown in green for saline (SS). Cocaine-induced fold changes from SS baseline shown on log2 scale for acute cocaine (SC), withdrawal from chronic cocaine (CS), and cocaine challenge following prolonged withdrawal (CC). Mono/di/trimethylation – me1/2/3; acetylation – ac; phosphorylation – ph; dark red circle, pval < 0.05; light red circle, pval < 0.1. (**B**) H2A.Z protein abundance in NAc across treatment groups as revealed by histone mass spectrometry. (**C**) mRNA expression of the *H2afz* gene encoding the H2A.Z variant. (**D**) Genomic distribution of H2A.Z according to ChIP-seq, which indicated primary enrichment of read counts over gene promoters. (**E**) Genome-browser view of the *Fosb* (upper panel) and *Fos* (lower panel) gene loci showing H2A.Z ChIP-seq tracks (H2A.Z signal normalized to total H3 ChIP-seq). (**F**) Heatmap shows MSN-subtype specific mRNA expression of genes that are selectively upregulated in D1 MSNs with cocaine challenge after prolonged withdrawal from chronic cocaine (CC), and not induced by acute cocaine in drug-naïve mice (SC). (**G**) Overlap of genes losing H2A.Z following withdrawal from chronic cocaine (CS) and those genes that are sensitized to cocaine challenge in D1 MSNs (shown in Fig. 3F), shows that most of these relapse-primed are marked by H2A.Z loss on withdrawal.

To investigate withdrawal-related changes in H2A.Z abundance genome-wide, we performed ChIP-seq for H2A.Z and, simultaneously, for total histone H3, which was used for chromatin normalization (Fig 3D-G). At baseline (SS), genomic H2A.Z binding in the NAc was dramatically enriched over promoter elements and flanked TSSs (Fig. 3D), corroborating recent reports for H2A.Z distribution in mouse hippocampus (*45*). Strikingly, following prolonged cocaine withdrawal, H2A.Z enrichment was markedly lowered at promoters and NAc-specific enhancer regions, including enhancer elements upstream of *Fosb* as just one specific example (Fig. 3E, upper panel). In contrast, depletion over *Fos* was less pronounced (Fig. 3E, lower panel).

Loss of H2A.Z was recently shown to increase the accessibility of transcription factor binding sites, predominantly for the AP-1 FOS and JUN families, in cultured human cells (*46*). These observations suggest that H2A.Z blocks promiscuous transcription factor access, and that withdrawal-associated H2A.Z depletion might sensitize genes for induction upon cocaine relapse. Hence, we examined the cohort of genes that are significantly upregulated in D1 MSNs upon cocaine relapse after withdrawal but that are not induced by acute cocaine in drug-naïve animals (Fig. 3F). We found that these relapse-primed genes, which remain unchanged during withdrawal, are characterized by withdrawal-related loss of H2A.Z binding (Fig. 3G), and regulate processes that govern synaptic plasticity, trans-synaptic signaling, and neuron projection morphogenesis (fig. S11A). Computational IPA revealed that these genes are targeted by transcription factors that are well-known to be activated by cocaine in D1 MSNs, including CREB1, C/EBP, and EGR1 (fig. S11B) (*12, 20, 47–49*).

### H2A.Z-specific histone chaperone ANP32E supports cocaine reward processing

The histone chaperone ANP32E (acidic leucine-rich nuclear phosphoprotein 32 family member E) was recently identified as a highly specific H2A.Z chaperone that removes, but does not deposit, H2A.Z from nucleosomes (*29*). We found that levels of ANP32E associated with chromatin in NAc is increased after cocaine withdrawal (unpaired T test p = 0.009, Fig. 4B), implicating a role for this protein in cocaine-induced H2A.Z eviction and downstream transcriptional regulation. To explore whether ANP32E is required for cocaine-induced depletion of H2A.Z, we generated an adeno-associated virus (AAV1) vector containing Cre-dependent short hairpin RNAs to knockdown (KD) *Anp32e* (shAnp32e) in D1 MSNs of D1-Cre mice. Intracranial injection of shAnp32e into the NAc lead to effective KD of ANP32E in this region (Fig. 4C). Additionally, we found that the ability of chronic cocaine exposure to deplete H2A.Z from NAc chromatin was abolished upon ANP32E KD (Fig. 4C), thus directly linking cocaine regulation of ANP32E in D1 MSNs to the cocaine-induced depletion of H2A.Z in this brain region.

**Fig. 4.**
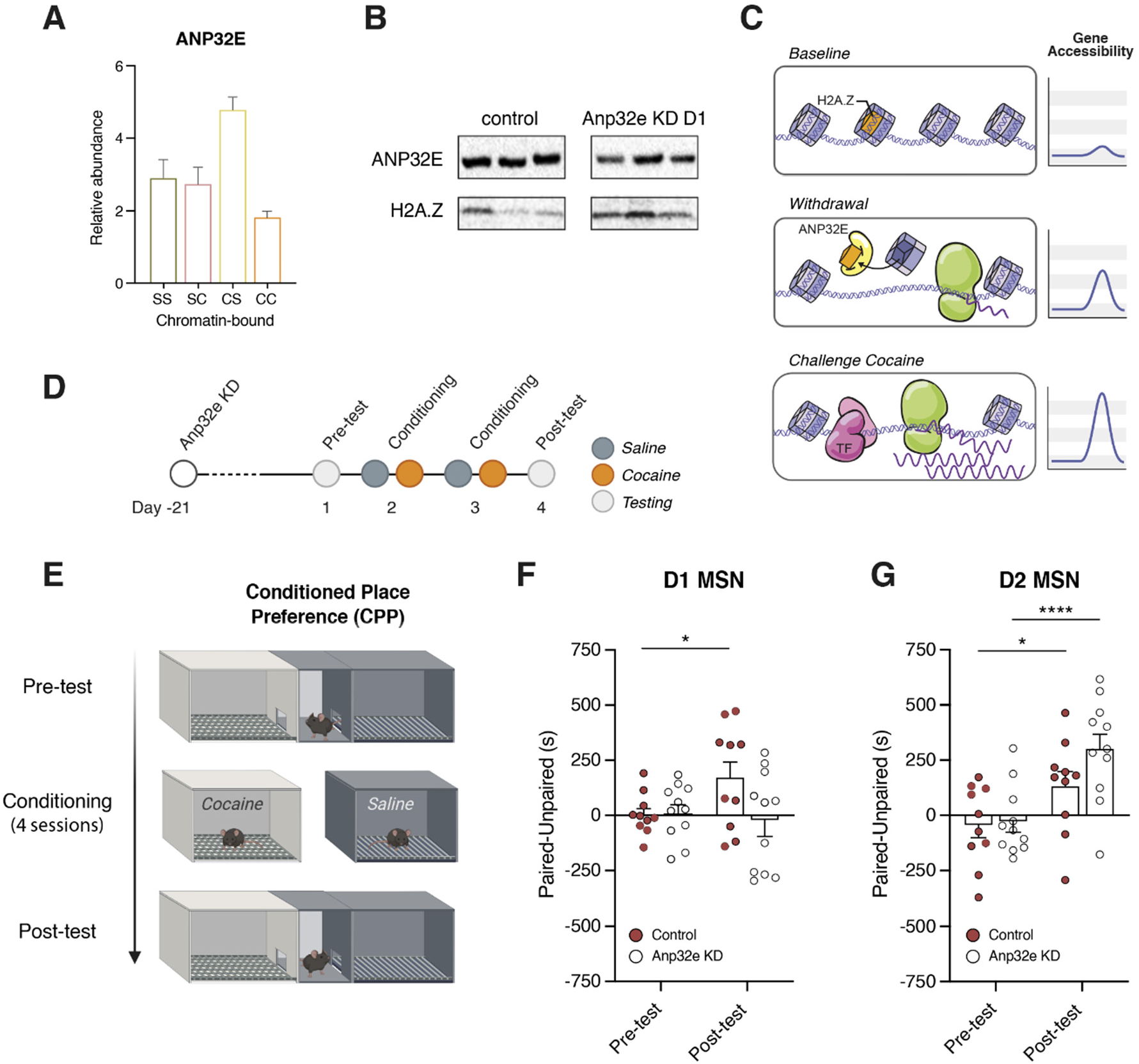
The H2A.Z-specific histone chaperone ANP32E regulates rewarding responses to cocaine. (**A**) Mass spectrometry of NAc chromatin showed that ANP32E abundance is increased with prolonged withdrawal from chronic cocaine (CS) (unpaired T test, p = 0.009, N = 6 per group). (**B**) Western blot revealed that *Anp32e* knockdown (KD) in D1 MSNs prevents H2A.Z depletion in the NAc following chronic cocaine exposure (6 days of cocaine following D1 MSN *Anp32e* KD using Cre-dependent shAnp32e AAV delivered to the NAc). (**C**) Model. Acute cocaine in drug-naïve mice increases H2A.Z acetylation with immediate-early gene expression. Following repeated cocaine, H2A.Z is evicted from chromatin concurrent with increased ANP32E abundance, linked to heightened gene responses upon cocaine challenge. (**D**) Experimental timeline for cocaine induced CPP in mice with a D1 MSN- or D2-MSN-specific *Anp32e* KD. (**E**) Schematic of cocaine induced CPP. (**F**) Preference scores for the cocaine-paired chamber in control mice and mice with D1-MSN-specific KD of *Anp32e* in the NAc (*F*(_1,19_) = *p* = 0.04, confirmed by Holm-Šídák’s multiple comparisons test). Data are mean ± s.e.m.; p values by Wilcoxon matched-pairs signed-rank test. (**G**) Preference scores for the cocaine-paired chamber in control mice and mice with D2-MSN-specific KD of *Anp32e* in the NAc (*F*(_19,19_) = 2.804, *p* = 0.015). Data are mean ± s.e.m.; p values by Wilcoxon matched-pairs signed-rank test.

To explore whether circuit-specific KD of *Anp32e* impacts cocaine-conditioned behaviors, we used an unbiased CPP paradigm, which provides an indirect measure of cocaine reward (Fig. 4D). As expected, *Anp32e* wild-type mice (shScr control) spent significantly more time in the compartment that was associated with cocaine experience in the post-test compared to their pre-test (*F*(_1,19_) = *p* = 0.04, confirmed by Holm-Šídák’s multiple comparisons test), indicating increased preference for the cocaine-paired environment in this group. In contrast, shAnp32e KD in D1-Cre mice did not form a preference for the chamber that was paired with cocaine experience in the post-test (Fig. 4F). These data indicate that ANP32E functions in D1 MSNs of NAc to facilitate cocaine-related reward processing. Notably, sex-specific effects of cocaine in the NAc have been recently identified as important factors enhancing motivation for cocaine in females (*50*). We thus investigated ANP32E function in female mice for drug-related behavior in the CPP paradigm and confirmed that D1-specific KD of *Anp32e* expression has a similar effect on cocaine-related learning as seen in males (*F*(_1,18_) = 8.017, *p* = 0.011; confirmed by Holm-Šídák’s multiple comparisons test; fig. S12).

We next tested whether ANP32E expression in D2 MSNs might also contribute to cocaine CPP, utilizing the Cre-dependent shAnp32e AAV1 in D2-Cre transgenic mice. In contrast to D1 circuit-selective *Anp32e* KD, which effectively blocked cocaine CPP, we found that the D2-specific knockdown of *Anp32e* enhanced cocaine-related reward learning in this behavioral paradigm (*F*(_19,19_) = 2.804, *p* = 0.015; confirmed by Holm-Šídák’s multiple comparisons test; Fig. 1G). These results collectively provide new insight into circuit-specific epigenetic and transcriptional priming by ANP32E employed by drugs of abuse to modify brain function and behavior.

## DISCUSSION

We showed previously that prolonged withdrawal from cocaine self-administration results in transcriptional priming of large sets of genes that are upregulated by cocaine relapse across the brain’s reward circuitry, with especially prominent effects seen in NAc (*42*). Predicted upstream regulators of this relapse-related gene program included AP-1 and CREB, which have long been implicated in the actions of drugs of abuse (*11, 13, 51*), along with several novel transcription factors such as E2F, which has since been validated as well (*43*). However, these analyses were based on whole-tissue transcriptome data, with still limited information available about the transcriptional effects of cocaine exposure, withdrawal, and relapse selectively in D1 vs. D2 MSNs, which display major, in some cases opposite, regulation by the drug (*2, 4, 10, 16, 52*).

Our cell-type-specific data show that acute cocaine and prolonged withdrawal predominantly impact gene regulation in D1 MSNs with more limited responses seen in D2 MSNs. Notably, D1 MSN chromatin accessibility is robustly increased in response to acute cocaine, this increase in sustained during prolonged withdrawal from chronic cocaine, and increased still further upon a relapse dose of the drug. Such increases in chromatin accessibility in D1 MSNs affect prominently both gene-regulatory enhancer and promoter elements of genes that regulate neuronal plasticity and metabolism. Interestingly, these findings are akin to a recently reported epigenetic priming mechanism in hippocampus during memory formation and recall, which also involves lasting and widespread ‘opening’ of chromatin and enhanced transcriptional responses, even though no change in respective baseline transcription was observed (*53, 54*).

In the present study, we found that cocaine-related changes in D1 MSN chromatin are linked to increases in reward-related gene expression upon drug relapse. Our bioinformatics analyses predict, again, that CREB and AP-1 transcription factors, among others, are upstream regulators of relapse-related and primed gene expression in D1 MSNs, thus validating prior work on the initial actions of drugs of abuse that implicates these factors in cocaine sensitization and cellular activity (*11, 13, 49, 55–57*), discussed above. These findings support our central hypothesis that cocaine regulation of the epigenetic landscape in NAc neurons determines the inducibility of discrete gene programs that characterize drug relapse. Our data directly establish that a histone variant, H2A.Z, plays a key role in this process.

This is the first study to implicate H2A.Z in the actions of cocaine or any other drug of abuse and underscores the value of utilizing unbiased experimental approaches to study the epigenetic basis of transcriptional regulation. Indeed, earlier ChIP-seq maps of cocaine regulation of several well-studied histone modifications in NAc revealed relatively limited overlap with changes in gene expression revealed by RNA-seq (*15*). By contrast, by following our open-ended analyses and identification of H2A.Z depletion as the most prominent histone change in NAc after prolonged withdrawal, we found that well more than 50% of genes in D1 MSNs that are primed for increased expression upon relapse exhibit H2A.Z depletion at their promoter or enhancer regions. Our proteomic analysis identified several additional histone modifications that are prominently regulated in NAc by cocaine withdrawal which now warrant similar characterization in future investigations.

To date, the functional relationship between H2A.Z and transcription remains uncertain, as H2A.Z has been associated with both positive and negative effects on gene expression, with its acetylation having a positive impact (*46, 58, 59*). Notably, the removal of H2A.Z from chromatin was shown to be critically important for stimulus- and training-induced gene expression in hippocampus (*58*), and hippocampal H2A.Z depletion resulted in enhanced memory formation. Furthermore, the loss of H2A.Z was recently found to cause a dramatic increase in the accessibility of AP-1 transcription factor binding sites in cultured cells (*46*), indicating that H2A.Z may prevent promiscuous transcription factor access and allows for rapid gene induction upon removal. Eviction of this variant is dependent upon ANP32E, an H2A.Z-specific histone chaperone (*29*). Our data are the first to show a function for this chaperone in NAc D1 and D2 MSNs in controlling rewarding responses to cocaine, with *Anp32e* expression (and H2A.Z depletion) in D1 MSNs supporting cocaine CPP, but limiting cocaine CPP in D2 MSNs.

Overall, histone variants and their chaperones have received limited attention in epigenetic studies of substance use disorders. Our data implicate H2A.Z histone variant exchange in the withdrawal-related epigenome remodeling that is linked to latent dysregulation of stimulus-induced gene expression upon drug relapse. Together, these findings present epigenetic priming involving ANP32E and H2A.Z depletion as a promising new clinical target that drugs of abuse engage in modifying brain function and behavior in lasting ways.

## METHODS

### Animals

Adult male C57BL/6 mice (8-16 weeks) were housed five per cage on a 12-h light-dark cycle at a constant temperature (23°C) and had free access to food and water ad libitum. They were assigned randomly to two groups treated for 10 consecutive days with either saline or cocaine (20 mg/kg) intraperitoneal injections and then subjected to a 30-day withdrawal period. Afterwards, the two groups were subdivided into another two groups receiving single injections of either saline or cocaine 1 h before the NAc tissue was harvested to construct a total of four subgroup conditions: control (SS, Sal-Sal), acute cocaine (SC, Sal-Coc), withdrawal (CS, Coc-Sal), and cocaine challenge (CC, Coc-Coc). For stereotactic surgeries, mice were anesthetized with ketamine (100 mg/kg) and xylazine (10 mg/kg). Syringe needles (33G, Hamilton) were used to bilaterally infuse 1 µl of virus at a 0.1 µl/min flow rate in head-fixed animals. Coordinates for NAc from Bregma: AP + 1.6 mm, ML + 1.6 mm, DV – 4.5 mm, 10° angle. All mice handling was undertaken in accordance with the Institutional Animal Care and Use Committee guidelines at Icahn School of Medicine at Mount Sinai.

### Western blots

Frozen NAc tissue samples were homogenized and incubated with agitation at 4°C for 30 min in RIPA lysis buffer (10 mM Trizma Base, 150 mM NaCl, 1 mM EDTA, 0.1% SDS, 1% Triton-X-100, 1% sodium deoxycholate, pH 7.4, protease inhibitors). Protein concentrations were determined by Pierce BCA protein assay (Thermo Scientific, #23225) and were normalized across samples using RIPA lysis buffer. Protein sample volumes were prepared accordingly to contain 25% 4X Laemmli Sample Buffer (Bio-Rad, #1610747) and 10% 2-Mercaptoethanol (Bio-Rad, #1610710) and were heated to 95°C for 5 min before being separated by SDS-PAGE with Criterion Precast Gels (Bio-Rad, 4–15% Tris/Glycine) and then transferred onto 0.2 µm nitrocellulose membranes (Bio-Rad, Trans-Blot, midi format). The membranes were blocked in 5% BSA in TBS supplemented with 0.1% Tween 20 (TBST) at RT for 1h. Primary antibodies were diluted 1:1000 in 2.5% BSA in TBST and incubated overnight at 4°C. Primary antibodies are listed below. The membranes were washed in TBST thrice for 10 min each, followed by incubation with HRP-conjugated secondary antibodies diluted 1:10000 in 2.5% BSA in TBST at RT for 2 h. The membranes were washed again three times and imaged with a Fujifilm LAS-4000 imager. Primary antibodies used were anti-H2AZ (GeneTex, GTX108273), anti-ANP32E (OriGene, TA351339), anti-H3 (Abcam, ab1791), and anti-H2AZ (Active Motif, #39013).

### Conditioned place preference (CPP)

An unbiased CPP test was carried out using three chambered CPP Med Associates boxes and software wherein two end chambers have distinct visual (gray vs. striped walls) and tactile (small grid vs. large grid flooring) cues to allow differentiation. All sessions were carried out in a dark room with ambient temperature. On the pre-test session, the mice were allowed to explore all three chambers unencumbered for 20 min. Groups were adjusted to counterbalance any pre-existing chamber bias. The conditioning was carried out by pairing an injection of saline with one chamber in the morning and a second injection of cocaine (10 mg/kg) with the other chamber in the afternoon for two consecutive days. Each conditioning session lasted 45 min. CPP testing was carried out on the fourth day where each mouse was allowed to explore all three chambers freely for 20 min. Place preference score was taken as time on the cocaine-paired side – time on the saline-paired side.

### Subcellular protein fractionation

NAc tissue punches were fractionated into cytoplasmic, membrane, soluble nuclear, and chromatin-bound nuclear extracts using the Subcellular Protein Fractionation Kit for Tissues (Thermo Scientific, #87790). Cytoplasmic extraction buffer (CEB) which selectively permeabilizes the cell membrane and releases soluble cytoplasmic contents was added to the NAc tissues and transferred to Dounce tissue homogenizers (setting B). The tissues were homogenized for 20 strokes and then centrifuged at 500xg for 5 min into new tubes with the Pierce Tissue Strainer that removes tissue debris. The supernatants (cytoplasmic extracts) were immediately transferred to clean tubes. The remaining pellets were then mixed and vortexed at maximum setting for 5 sec with the membrane extraction buffer (MEB), which dissolves plasma, mitochondria, and ER/Golgi membranes without solubilizing the nuclear membrane. The mixtures were then centrifuged at 3000xg for 5 min. The supernatants (membrane extracts) were again transferred to new tubes. The pellets containing intact nuclei were vortexed at maximum setting for 15 sec with the nuclear extraction buffer (NEB) and then incubated at 4°C for 30 min with gentle mixing. They were centrifuged at 5000xg for 5 min which yielded the supernatants with soluble nuclear extracts. Lastly, the chromatin-bound extraction buffer was prepared by adding 5 µl of 100 mM CaCl_2_ and 3 µl of micrococcal nuclease to 100 µl of room temperature NEB and then added to the pellet and vortexed again. The tubes are incubated in a 37°C-water bath for 15 min followed by vortexing again and centrifuging at 16000xg for 15 min. The supernatant (chromatin-bound nuclear extract) was then transferred into new tubes. All extraction buffers contained protease inhibitors (c0mplete mini, EDTA-free #40774700) and were kept on ice until use.

### Chromatin immunoprecipitation followed sequencing (ChIP-seq)

Chromatin was extracted using truChIP Chromatin Shearing Kit with formaldehyde and prepared according to manufacturer’s Low Cell protocol (Covaris PN010179 Rev O, Dec 2020). 11.1% formaldehyde was made fresh from 16% concentrated formaldehyde. Room temperature (RT) 1X Fixing Buffer A was added to samples and the 11.1% formaldehyde was then added to make a final concentration of 1% only. Samples were incubated for 5 min on rocker at RT to allow for efficient cross-linking and terminated immediately by adding quenching Buffer E and incubating for another 5 min. Supernatants were discarded and the tissue samples were washed with cold 1X PBS twice. Lysis Buffer B was added and samples were homogenized using the Dounce Tissue homogenizer (∼20 strokes) and left to incubate on rocker for 10 min at 4°C. Samples were then centrifuged at 1700xg for 5 min at 4°C and the supernatant discarded followed by resuspension in wash buffer C and incubated for 10 min at 4°C. Supernatants were discarded after centrifuging at 2000xg for 5 min at 4°C and shearing buffer D3 was added without disturbing the nuclei pellet followed by two rounds of centrifugation at 1700xg for 5 min at 4°C while the supernatants were decanted. The nuclei pellets were resuspended in shearing buffer D3 and transferred to AFA microtubes (8 microTUBE-130 AFA Fiber H Slit Strip V2, PN520239) for low cell chromatin shearing using Covaris ultrasonicator with the following settings: 75.0 peak power; 15.0 Duty % Factor; 1000 Cycles; 11.3 Avg Power for durations of 810.0 sec and 540.0 sec, respectively.

After shearing, samples were transferred into pre-chilled microcentrifuge tubes and diluted 1:1 with Covaris 2X IP Dilution buffer and microcentrifuged at 10000xg for 5 min at 4°C before proceeding to IP with the supernatant. 30 µL of beads (Invitrogen Dynabeads Protein G) per IP was aliquoted into 1.5 ml Eppendorf tubes and separated on magnet for ∼1 min and the supernatant removed by aspiration. The beads were resuspended and mixed in 1 ml Block solution (1X PBS, 0.5% BSA) by inversion. Supernatant was separated on magnet and aspirated again and the wash process repeated once more. Finally, the beads were resuspended in 250 ml Block solution and 4 µg/per IP of H2A.Z and H3 antibodies, respectively, and the tubes rotated at 4°C for 2 h. The beads were collected on magnet and washed in 1 ml Block solution for a total of 3 washes before resuspension in 50 µl/per IP Block solution. Chromatin lysates from each sample were aliquoted respectively for H2A.Z and H3 antibody IP into 1.5 ml lo-bind protein Eppendorf tubes, 50 µl of the beads and 300 µl of block solution added and mixed by inversion. The IP tubes were rotated at 4°C overnight. The IP/bead mixture were transferred to new pre-chilled tubes and the beads were collected on magnet and the supernatant aspirated. 1 ml of cold RIPA wash buffer (50 mM HEPES-KOH pH 7.5, 500 mM LiCl, 1 mM EDTA, 1% NP-40, 0.7% Na-Deoxycholate) was added and mixed by inversion. The beads were collected, followed by a total of 5 RIPA washes before resuspension in 1 ml Final ChIP Wash buffer (1X TE, 50 mM NaCl) and subsequent collection on a magnet. The beads were then resuspended in 210 µl Elution buffer (50 mM Tris-HCl pH 8.0, 10 mM EDTA, 1% SDS) and incubated at 65°C for 30 min with 400 rpm on a thermomixer. The beads were collected again on magnet and 200 µl of the supernatant transferred to fresh tubes. The tubes were then boiled at 95°C for 7 min for DNA reverse cross-linking. The DNA were purified using Qiagen MinElute PCR purification kit and eluted with 20 µl Buffer EB (10mM Tris-Cl, pH 8.5) and transferred to 8-strip PCR tubes.

The library was prepared according to NEBNext Ultra II DNA Library Prep Kit for Illumina (NEB#E7645L) protocol. The size distribution of the DNA libraries was validated using the Agilent Bioanalyzer High Sensitivity DNA chip, after which all libraries were sequenced with Genewiz/Azenta on an Illumina NovaSeq S4 machine using a 2 x 150 bp pair-end read configuration to a minimum depth of 30 million reads per sample. Raw reads were processed by trimming the adapter sequences using TrimGalore. Trimmed reads were aligned to the reference genome mm10 using HISAT2. Duplicate fragments were discarded, leaving only high-quality unique reads. Reads were sorted by chromosomal coordinates using samtools (*60*). MACS was utilized to obtain sites of H2A.Z (peaks) for each of the 4 conditions - SC, SS, CS and CC.

### NAc nuclei isolation for fluorescence-activated nuclei sorting (FANS)

Nuclear samples were obtained from frozen tissue, collected by bilateral NAc punch dissections from 1 mm-thick coronal brain sections using a 14G needle. Virally infected tissue was harvested under fluorescent light. Frozen NAc samples were mechanically dissociated and nuclei lysed using a glass douncer in ice-cold lysis buffer (10.94% w/v sucrose, 5 mM CaCl_2_, 3 mM Mg(CH_3_COO)_2_, 0.1 mM EDTA, 10 mM Tris-HCl pH 8, 1 mM DTT, in H_2_O). Homogenates were filtered through a 40 µm cell strainer (Pluriselect) into ultracentrifuge tubes (Beckman Coulter), underlaid with 5 mL of high-sucrose solution (60% w/v sucrose, 3 mM Mg(CH_3_COO)_2_, 10 mM Tris-HCl pH 8, 1 mM DTT, in H_2_O). The sucrose gradient was centrifuged at 24,400 rcf for 1 h at 4°C in a SW41Ti Swinging-Bucket Rotor (Beckman Coulter), and nuclei at the bottom of the gradient were resuspended in PBS. DAPI was added at a concentration of 0.5 μg/mL. Whole cell samples were obtained from fresh tissue, which was rotated at 37 °C for 45 min in 1 mg/mL papain suspended in digestion buffer (5% w/v D-trehalose, 0.05 mM APV, 0.0125 mg/mL DNAse, in Hibernate^TM^-A (Thermofisher, A1247501)). Tissue was then placed in FANS buffer (0.58 mg/mL albumin inhibitor (Worthington Biochemical, LK003182), 5% w/v D-trehalose, 0.05 mM APV, 0.0125 mg/mL DNAse, in Hibernate^TM^-A) and triturated using progressively smaller pipette tips. Samples were passed through a 70 µm filter and placed on top of a layer of 10 mg/mL ovomucoid-albumin in FANS buffer. The pellet was resuspended in FANS buffer and DAPI was added at a concentration of 0.5 μg/mL.

### Assay for transposase accessible chromatin with sequencing (ATAC-seq)

A published protocol was used for ATAC-seq (*61*). Nuclei pellets were resuspended gently in transposition reaction mix (25 µL 2X TD Buffer [Illumina Cat #FC-121-1030], 2.5 µL transposase [Illumina Cat #FC-121-1030] and 22.5 µL nuclease-free water) and incubated at 37°C for 30 min followed immediately with purification using Qiagen MinElute kit whereby transposed DNA were eluted in 10 µl Elution Buffer. Transposed DNA fragments were amplified initially using 1X NEBnext PCR master mix (New England Labs Cat #M0541), 25 µM customized Nextera PCR primers 1 and 2 and 0.6% final concentration of 100X SYBR Green I (Invitrogen Cat #S-7563) in a thermocycler with the following parameters: 72°C for 5 min; 98°C for 30 sec; 5 cycles of 98°C for 10 sec, 63°C for 30 sec; and 72°C for 1 min. A side qPCR amplification using a small aliquot of the amplified DNA was used to determine the necessary additional number of cycles to stop amplification prior to saturation (1/4 of max fluorescent intensity) to reduce GC and size bias. The remaining DNA were amplified to the correct number of cycles and then purified and eluted using Qiagen PCR Cleanup kit in 20 µl of Elution buffer. Library quality control (QC) was done using Agilent Bioanalyzer High Sensitivity DNA chip with the samples diluted 1:5. Raw reads were processed by trimming the adapter sequences using Cutadapt (*62*). Trimmed reads were mapped to the reference mm10 genome using HISAT. Duplicate fragments were discarded, retaining only unique reads. Reads were sorted by chromosomal coordinates using samtools (*60*). ngsplot tool was utilized to generate average profile plots for each sample (*63*).

### RNA-sequencing (RNA-seq)

RNA was extracted from frozen TRIzol LS (Ambion) homogenates using the Direct-zol RNA Microprep Kit (Zymo Research) following manufacturer instructions. Ribo-depleted sequencing libraries were prepared with the SMARTer Stranded Total RNA-Seq Kit v2 - Pico Input Mammalian (TaKaRa Biotech) following manufacturer instructions and sequenced with Genewiz/Azenta on an Illumina NovaSeq S4 machine using a 2 x 150 bp pair-end read configuration to a minimum depth of 30 million reads per sample. Differential expression was analyzed in R using the DESeq2. Significance cut-offs were of at least 30% expression fold change (|log2(FoldChange)| > log2(1.3)) and nominal p < 0.05. RRHO plots were generated using the RRHO2 package (github.com/RRHO2/RRHO2).

## Acknowledgements

Support for this work was provided by grants from the National Institutes of Health, DA047233 (EJN), DA007359 (EJN), AA027839 (PM) and from the Brain and Behavior Research Foundation (PM).

## EXTENDED DATA & FIGURE LEGENDS

**Extended Data Fig. 1.**
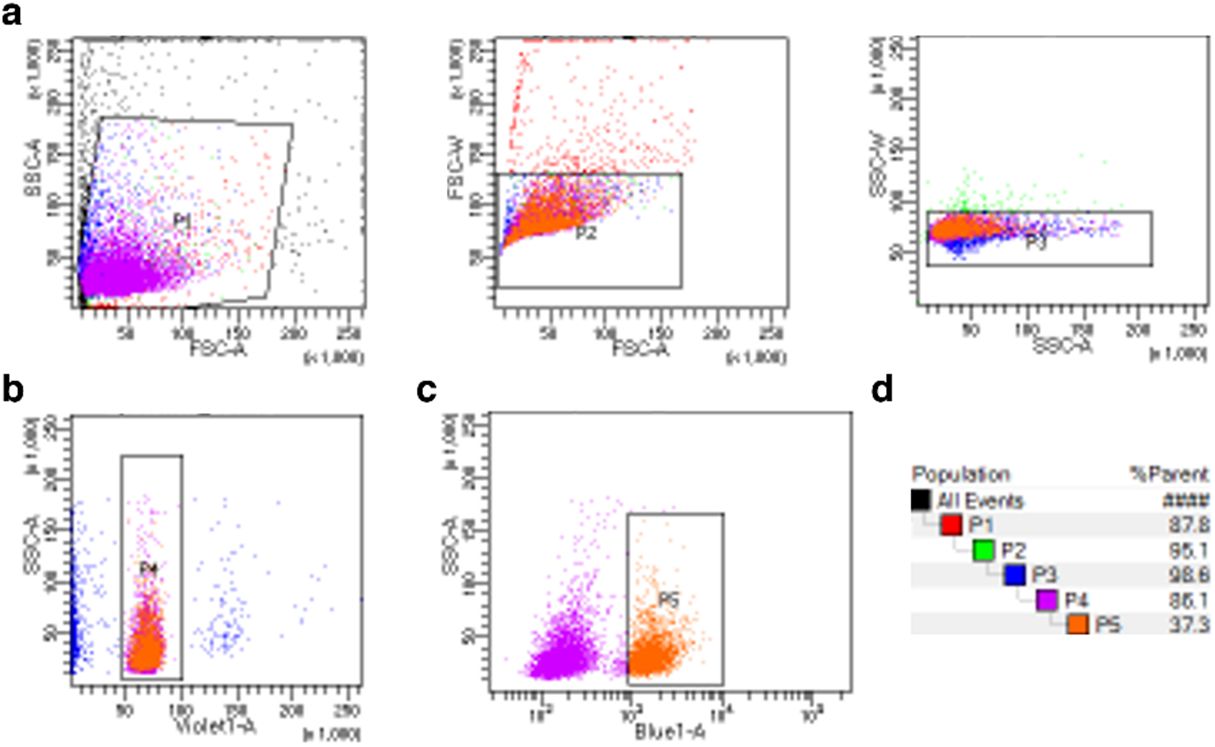
Fluorescence-activated nuclei sorting (FANS) to purify D1 and D2 MSN nuclei from NAc. Representative gating strategy for FANS. **(a)** First three gates on FSC-A vs. SSC-A (P1), FSC-A vs. FSC-W (P2), and SSC-A vs SSC-W (P3) retrieve nuclei as opposed to debris. **(b)** Gating strategy on Violet1-A vs. SSC-A (P4) retrieves single nuclei (DAPI+ events) as opposed to duplets. **(c)** The final gate FSC-A vs. Blue1-A (FITC channel) separates transgenically labeled nuclei (P5, GFP+ events) from wild-type nuclei. **(d)** Summary table of representative hierarchical gating strategy.

**Extended Data Fig. 2.**
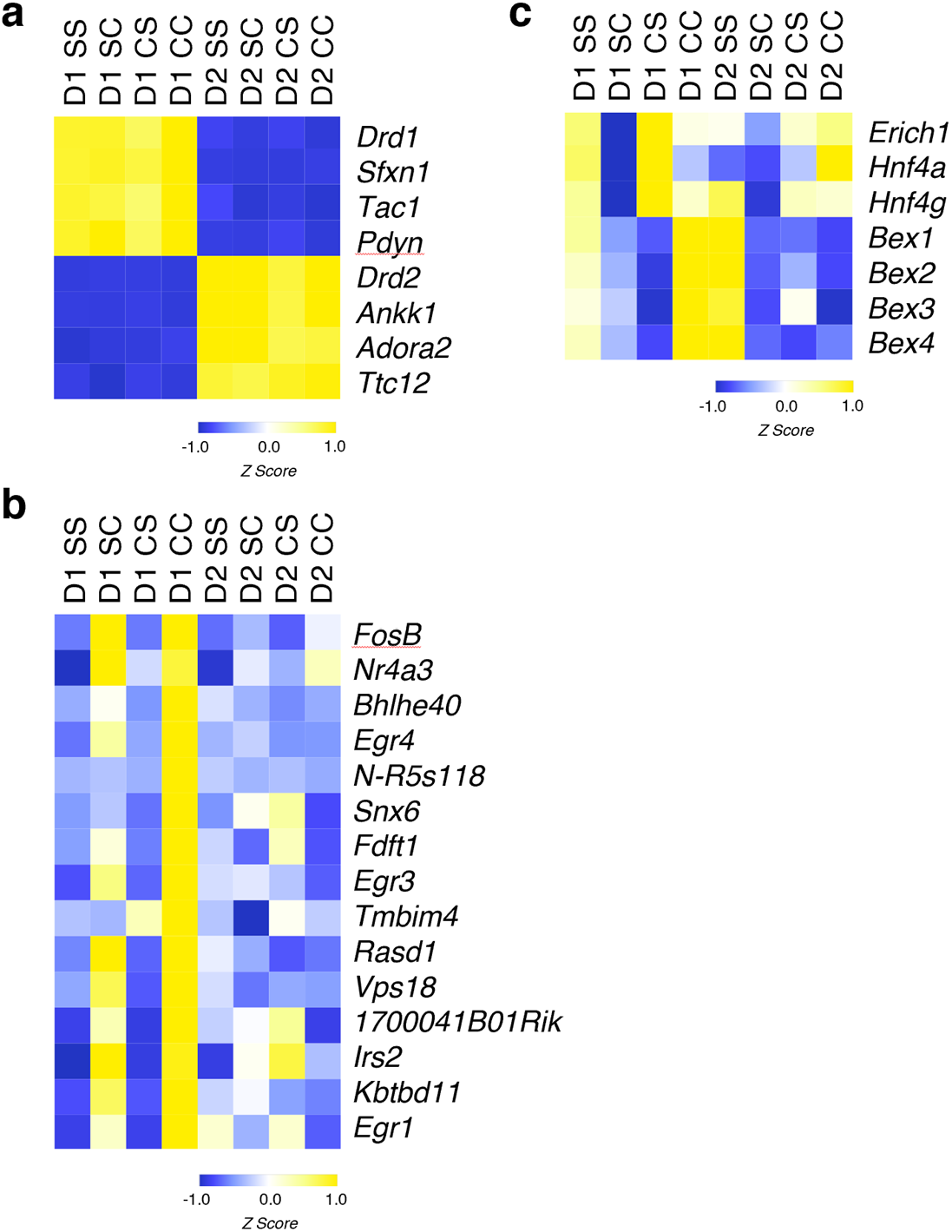
**(a)** Heatmap of mRNA expression (cpm) of well-characterized genes specific to D1 MSNs (*Drd1, Sfcn1, Tac1, Pdyn*) or to D2 MSNs (*Drd2, Ankk1, Adora2, Ttc12*), confirming successful sorting and purification of D1 and D2 MSN nuclei. **(b)** mRNA expression heatmap of top genes upregulated in D1 MSNs with cocaine challenge in withdrawal animals (D1 CC), indicating priming of cocaine-responsive genes when compared to acute cocaine (D1 SC). **(c)** mRNA expression heatmap of genes previously shown to be upregulated upon withdrawal from cocaine (*Erich1, Hnf4a, Hnf4g*) or depressed with cocaine or alcohol addiction in humans (Bex gene family, *Bex1-4*) (*43, 64*)

**Extended Data Fig. 3.**
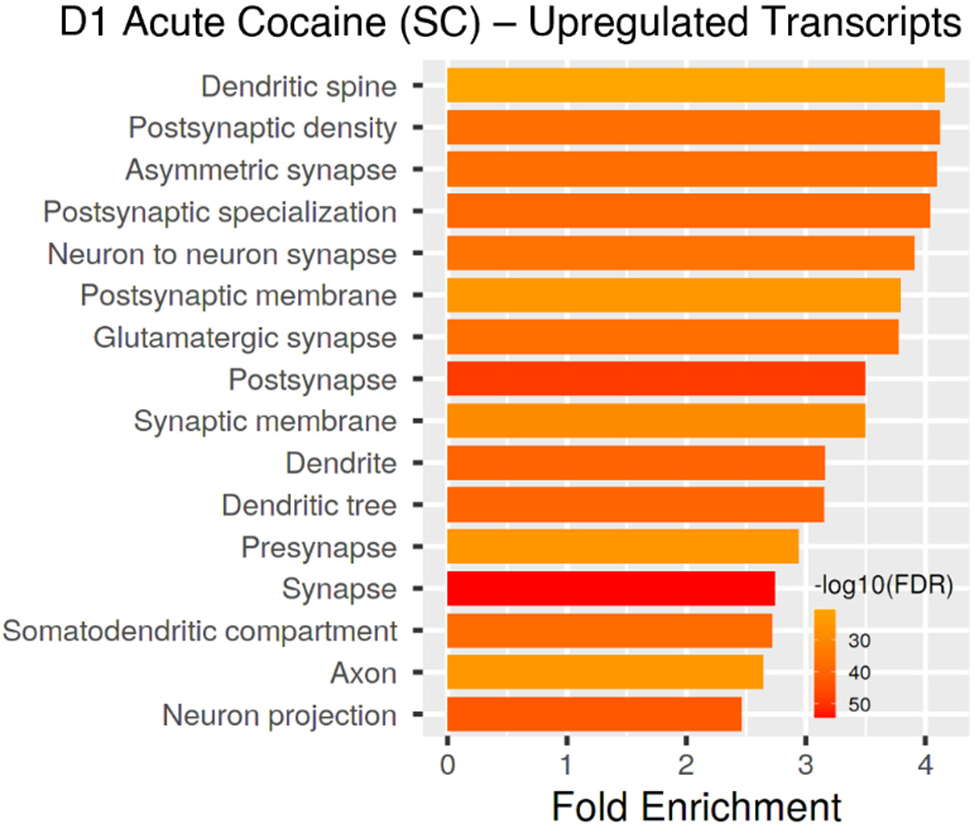
Gene ontology (GO) enrichment of genes that become upregulated with acute cocaine in D1 MSNs using ShinyGO 0.75 (2871 upregulated genes shown in top panel Fig. 1C).

**Extended Data Fig. 4.**
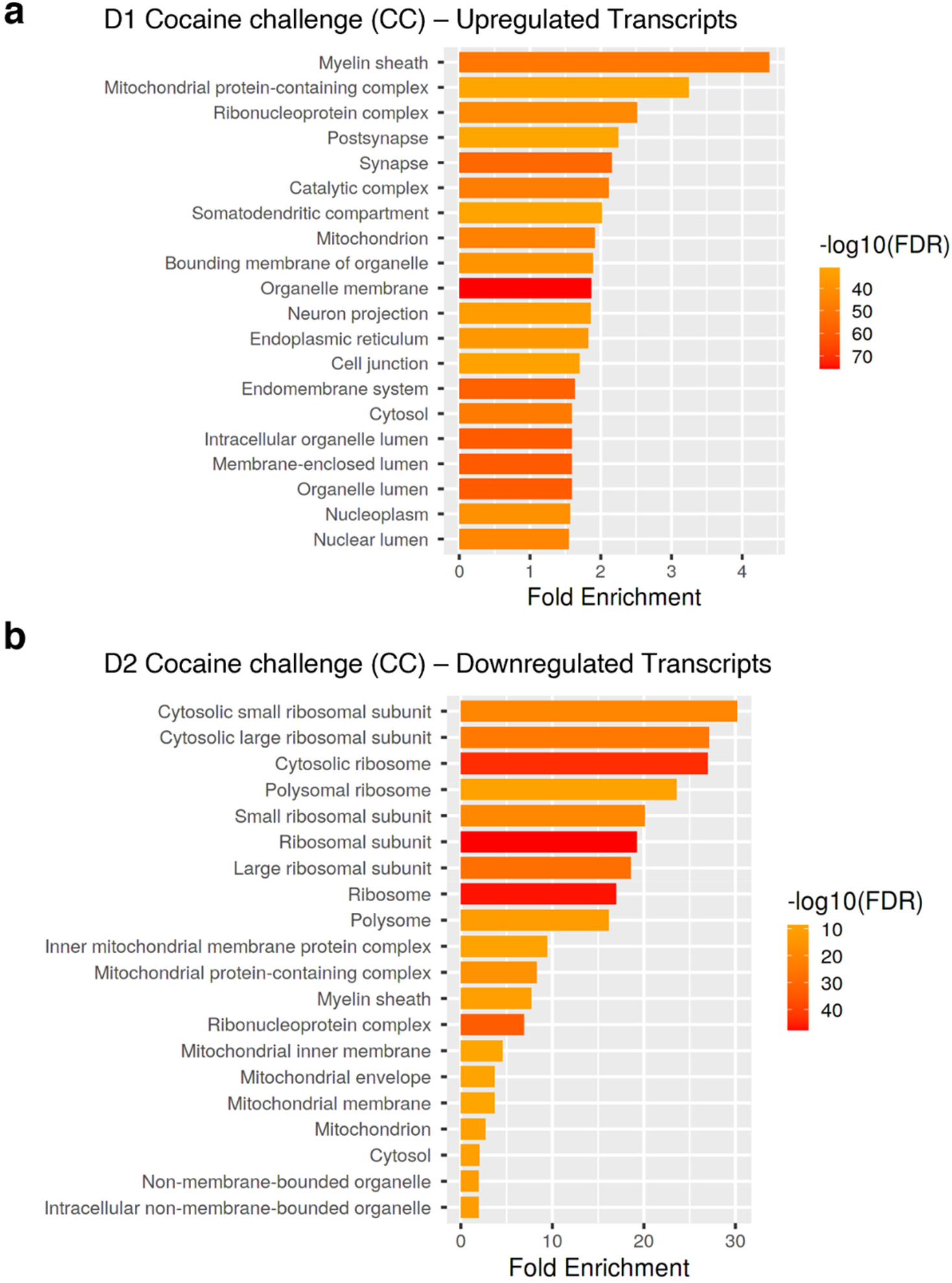
GO term enrichment analysis shows opposite gene regulation in D1 vs. D2 MSNs of NAc with cocaine challenge. **(a)** GO term enrichment of genes that are upregulated in D1 MSNs with cocaine challenge after prolonged withdrawal from chronic cocaine using ShinyGO 0.75 (4704 genes shown in lower right panel Fig. 1C). **(b)** GO term enrichment of genes that are downregulated in D2 MSNs with cocaine challenge after prolonged withdrawal from chronic cocaine using ShinyGO 0.75 (583 genes shown in lower right panel Fig. 1C).

**Extended Data Fig. 5.**
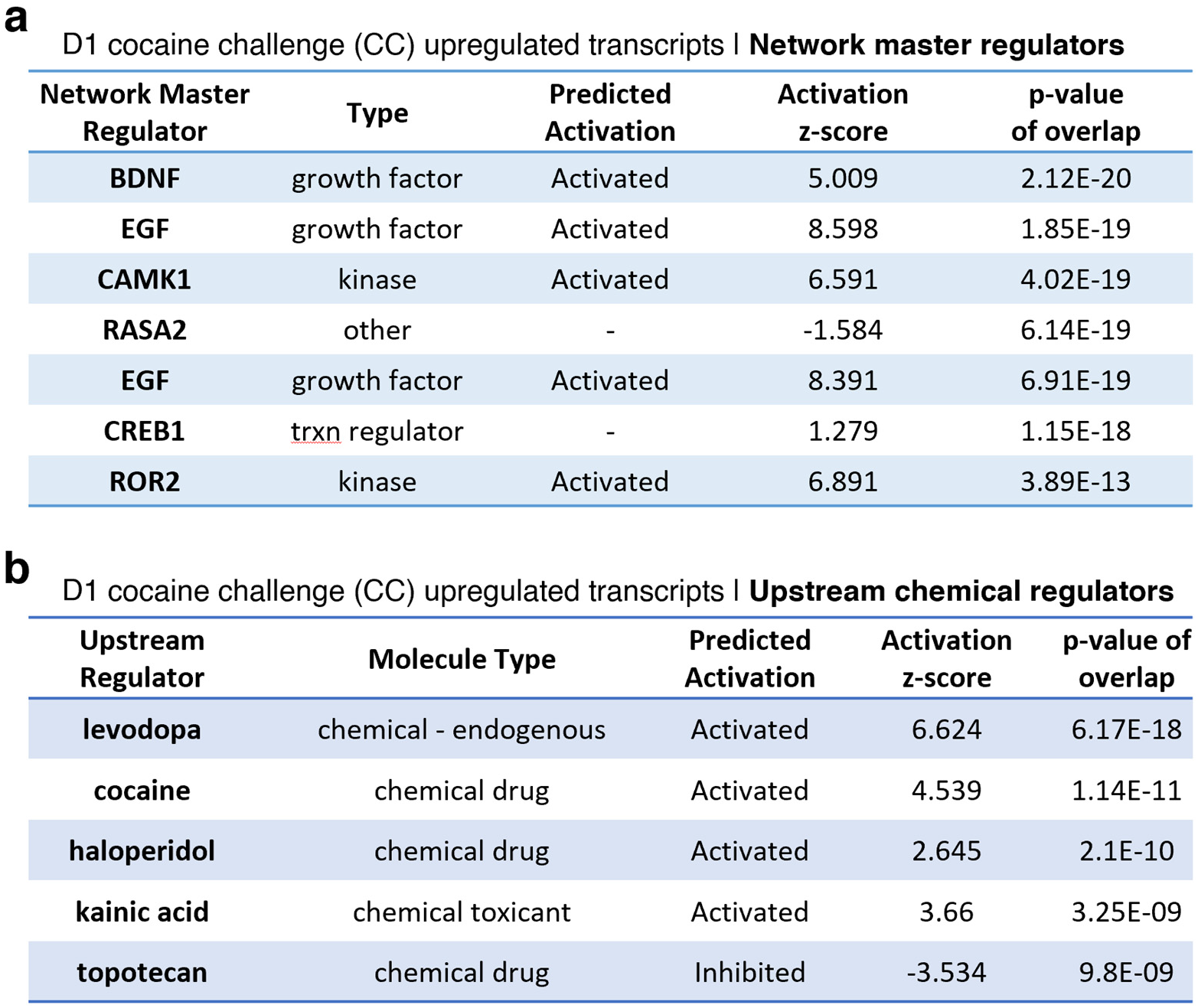
Upstream regulators of cocaine-induced gene programs in D1 MSNs of NAc. Network master regulators **(a)** and upstream chemical regulators **(b)** of gene programs that are induced by acute cocaine in D1 MSNs, as revealed by IPA (2871 upregulated genes shown in top panel Fig. 1C).

**Extended Data Fig. 6.**
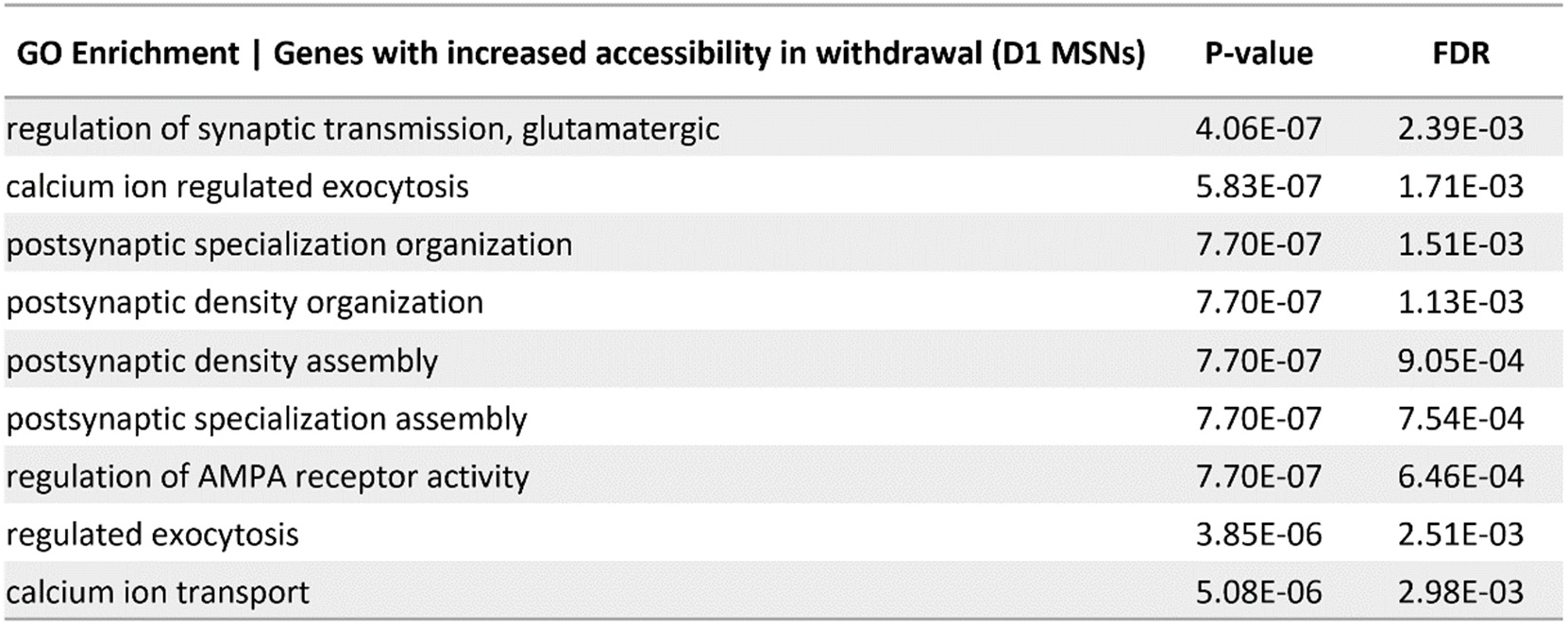
GO term enrichment analysis of genes that show increased accessibility in D1 MSNs following cocaine withdrawal (CS).

**Extended Data Fig. 7.**
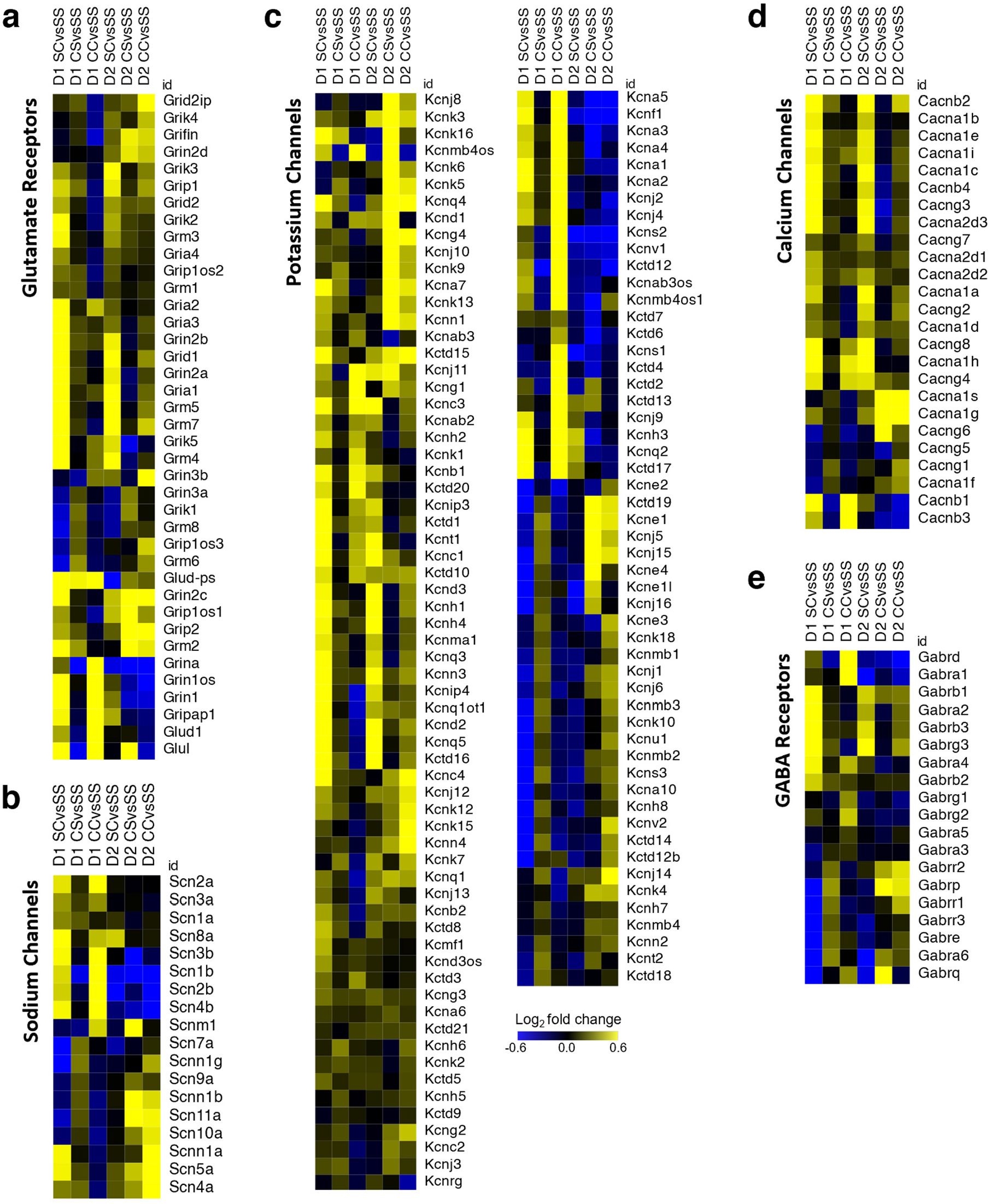
Excitability- and plasticity-related genes exhibit cell-type-specific regulation by cocaine. Heatmaps show mRNA gene expression of key ionotropic receptors and ion channels upon acute cocaine (SC), prolonged cocaine withdrawal (CS), and cocaine challenge after withdrawal (CC), in both D1 and D2 MSNs of NAc. These plasticity-related transcripts include **(a)** glutamate receptors, **(b)** sodium channels, **(c)** potassium channels, **(d)** calcium channels, and **(e)** GABA receptors.

**Extended Data Fig. 8.**
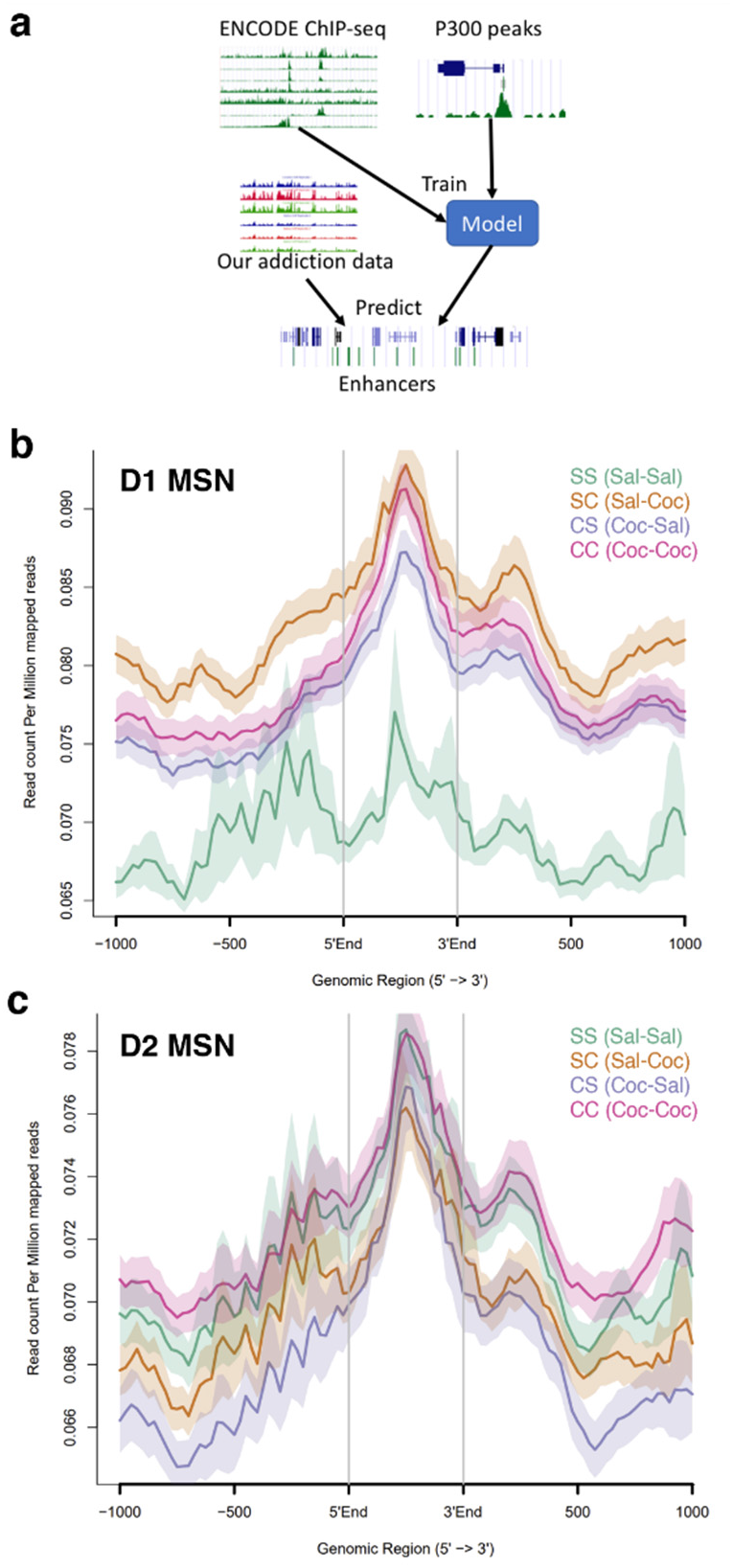
Identification of cocaine-regulated enhancers shows D1-specific accessibility changes. **(a)** Machine learning model, DeepRegFinder (*41*), identifies putative cocaine-regulated enhancer regions. DeepRegFinder utilizes 1-D convolutional and recurrent neural networks to extract features that represent characteristic chromatin modification patterns of representative enhancers (defined by P300) from ENCODE ChIP-seq data. The trained model was then used to identify enhancers on the whole genome. We used ChIP-seq data from a previous cocaine study to build a model to identify 7,796 cocaine-related enhancers in NAc for this study (*15*). Accessibility of these 7,796 putative enhancers shown for **(b)** D1 MSNs and **(c)** D2 MSNs in saline control animals (SS), after acute cocaine in drug-naïve mice (SC), after prolonged withdrawal from chronic cocaine (CS), and upon cocaine challenge in withdrawal animals (CC).

**Extended Data Fig. 9.**
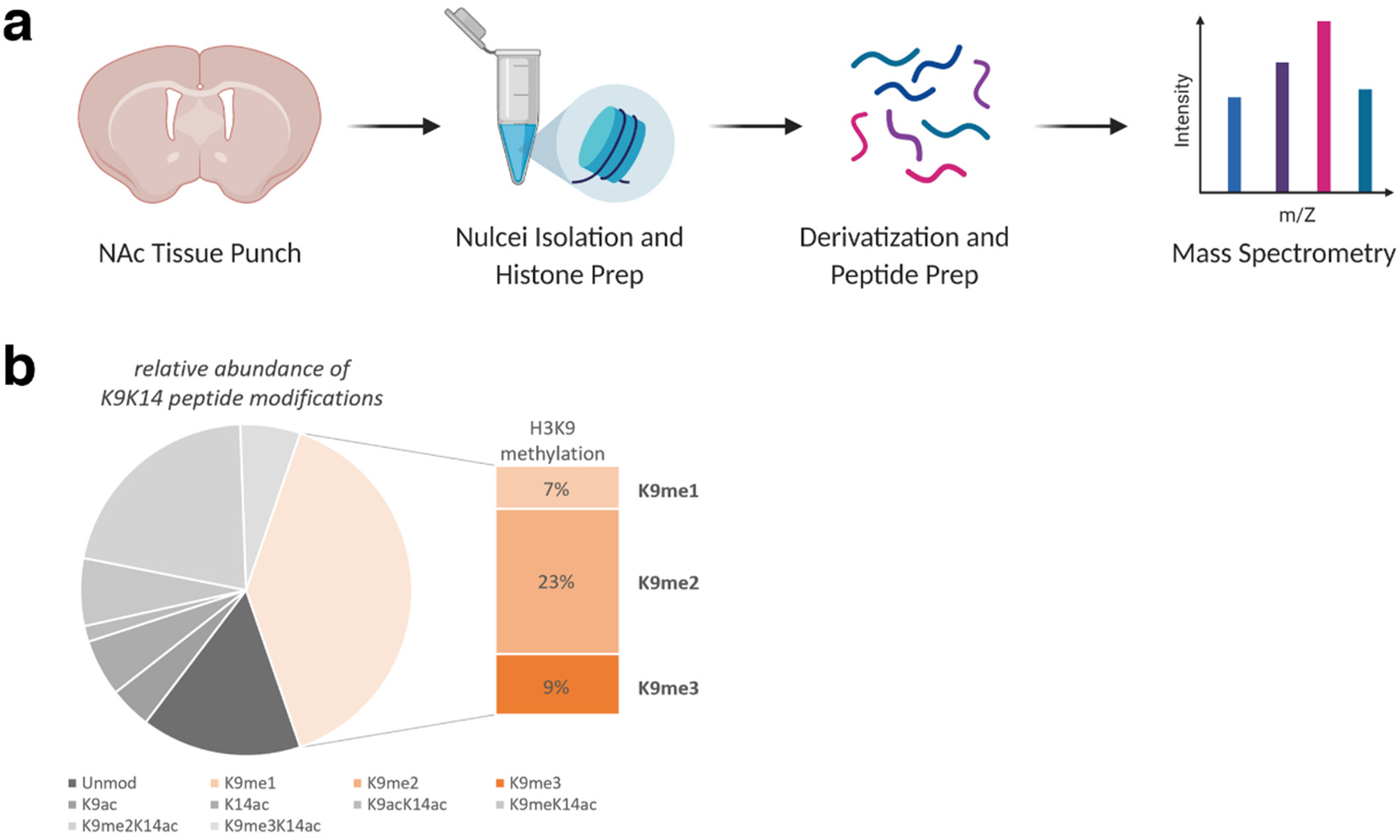
Mass spectrometry reveals changes in chromatin in NAc. **(a)** Experimental outline for histone mass spectrometry starting from NAc tissue punches, which is the starting material for nuclei isolation and histone protein extraction. The prepared histones are first derivatized using propionic anhydride to neutralize charge and block unmodified and monomethylated lysine residues, and are subsequently digested using trypsin, which, under these conditions, cleaves only arginine residues. The generated peptides are then analyzed using online LC-electrospray ionization-tandem mass spectrometry to identify modification sites. **(b)** Relative abundance of H3 lysine 9 (H3K9) and lysine 14 (H3K14) acetylation and methylation (me1/2/3 – mono/di/tri-methylation).

**Extended Data Fig. 10.**
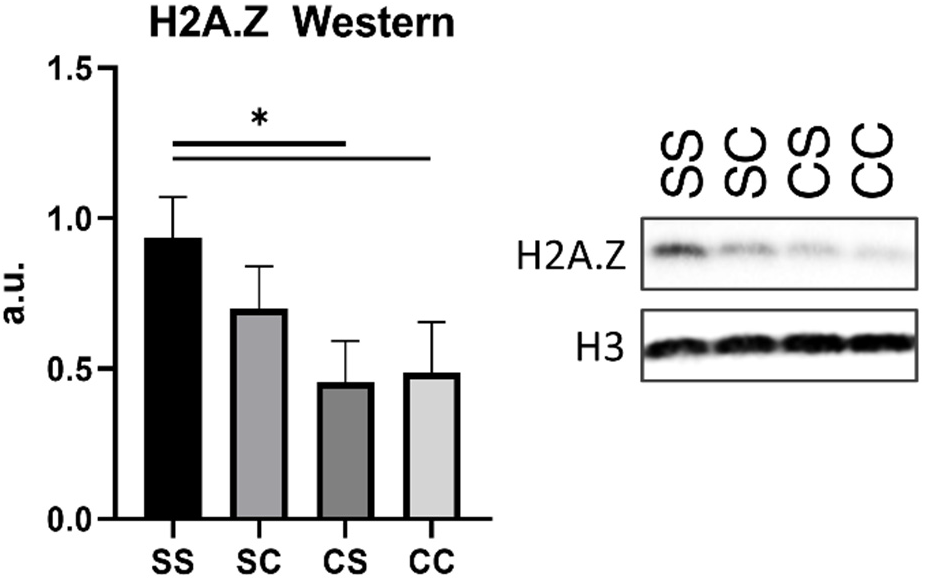
Western blot confirming mass spectrometry finding of depleted H2A.Z in NAc following prolonged withdrawal from chronic cocaine.

**Extended Data Fig. 11.**
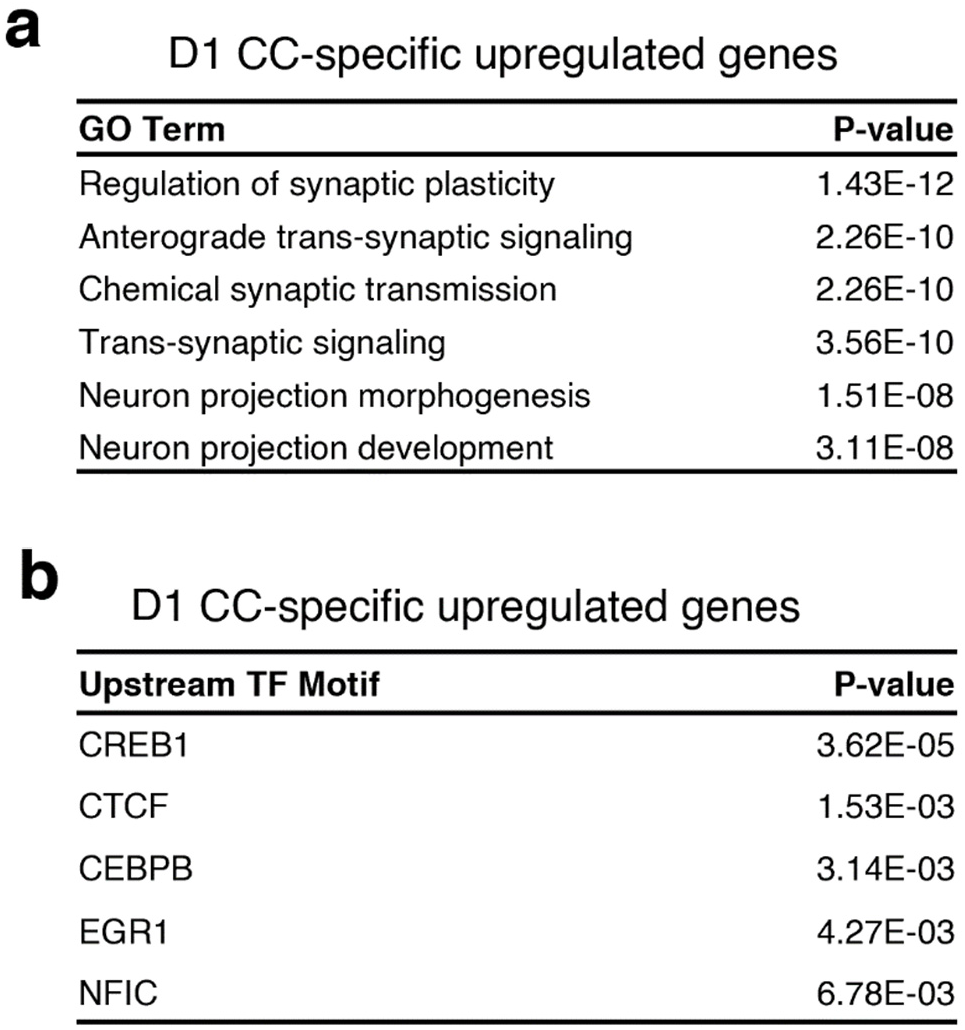
**(a)** GO term enrichment analysis of gene sets only induced with cocaine challenge in withdrawal, yet not with acute cocaine in drug-naïve mice. **(b)** GO term enrichment analysis of gene sets only induced with cocaine challenge in withdrawal, yet not with acute cocaine in drug-naïve mice.

**Extended Data Fig. 12.**
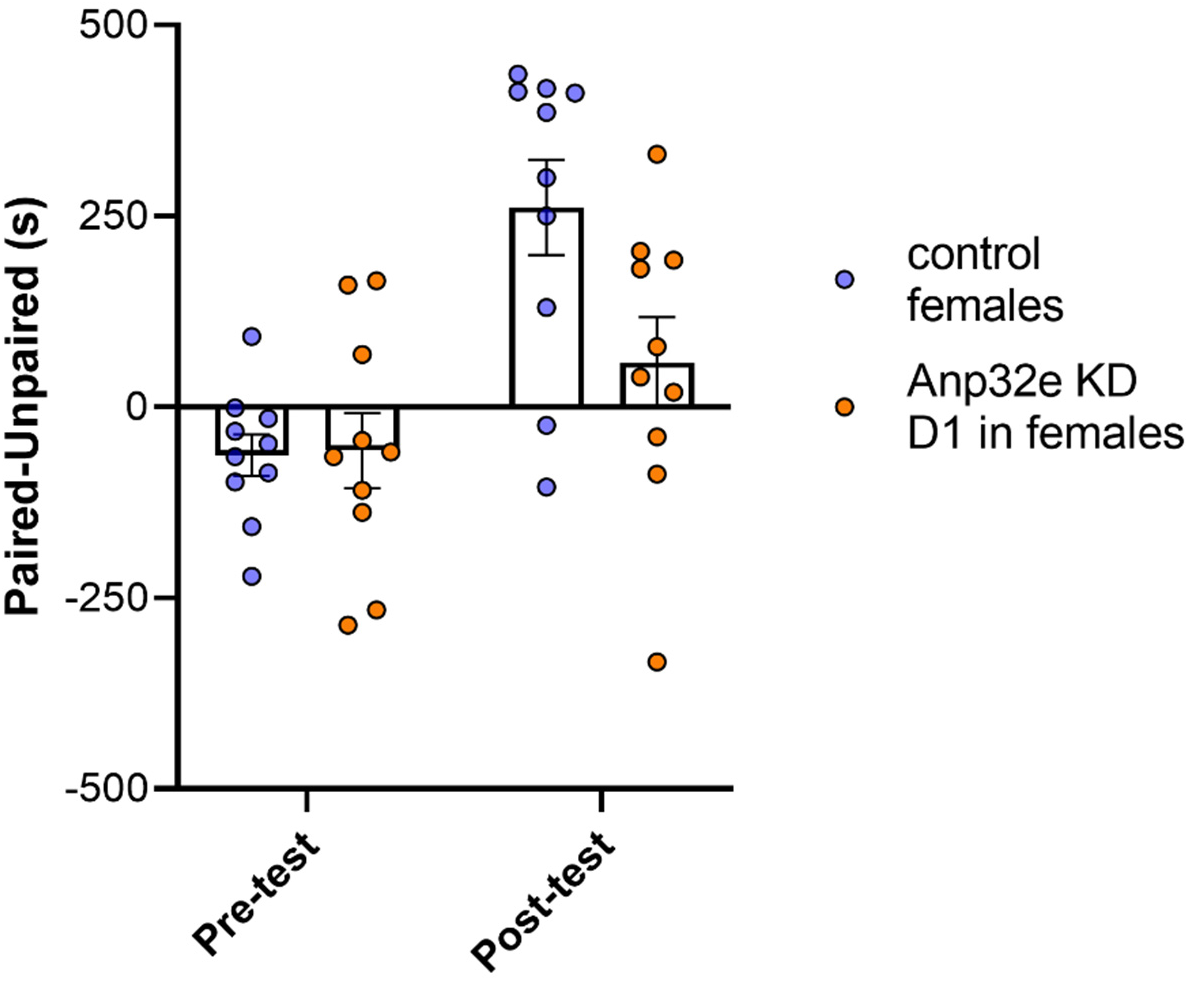
Cocaine CPP in female mice with cell-type-specific *Anp32e* KD in D1 MSNs. Preference scores for the cocaine-paired chamber in control mice and mice with subtype-specific KD of *Anp32e* in D1 MSNs of the NAc (*F*(_1,18_) = 8.017, *p* = 0.011; confirmed by Holm-Šídák’s multiple comparisons test). Data are mean ± s.e.m.; p values by Wilcoxon matched-pairs signed-rank test.

